# Dynamical independence reveals anaesthetic specific fragmentation of emergent structure in neural dynamics

**DOI:** 10.1101/2025.07.16.664881

**Authors:** Borjan Milinkovic, Anil K. Seth, Lionel Barnett, Olivia Carter, Thomas Andrillon

**Affiliations:** Melbourne School of Psychological Sciences. University of Melbourne, Melbourne, Australia; INSERM/Institut du Cerveau (ICM), Hôpital de la Pitié-Salpêtrière, Paris, France; Paris-Saclay Institut des Neurosciences, CNRS, Gif-sur-Yvette, Paris, France; Sussex Centre for Consciousness Science, Department of Informatics, University of Sussex, Brighton, UK; Canadian Institute for Advanced Research, Program on Brain, Mind, and Consciousness, Toronto, Canada; Monash Centre for Consciousness & Contemplative Studies, Monash University, Melbourne, Australia

**Keywords:** emergence, dynamical independence, macroscopic organisation, scale integration, consciousness, EEG, anaesthesia, higher-order brain dynamics, dimensionality reduction, multiscale structure

## Abstract

Conscious experience depends on the coordinated activity of neural processes that span multiple scales—from synapses to whole-brain dynamics. A recently introduced measure, *dynamical independence*, identifies, characterises and quantifies these multi-scale relationships using an information-theoretic dimensionality-reduction approach. Here, we use DI to examine changes in emergent dynamical structure in the human brain under three pharmacologically-distinct anaesthetic interventions (propofol, xenon, ketamine). Applied to source-reconstructed EEG, our analysis reveals that propofol and xenon, anaesthetics that abolish conscious report, exhibit more emergent but highly variable dynamic structure, indicating fragmented macroscopic dynamical organisation. By contrast, ketamine, which preserves dream-like phenomenology, shows the opposite pattern: reduced overall emergence yet a partial preservation of the macroscopic structure, mirroring wake. Further exploratory analyses revealed spatially localised source-level contributions to emergent dynamical structure, highlighting regional variations. Together, our results highlight drug-specific reconfigurations of emergent dynamical structure under anaesthesia, dissociate the *amount* of emergence from the *organisation* of emergent dynamics, and caution against equating emergence with level of consciousness.

## 1 Introduction

Transitions between global states of (un)consciousness have often been associated with changes of the structured dynamical interactions in neural activity across multiple spatial and temporal scales [1–6]. In neuroscience, these spatial scales are broadly and somewhat arbitrarily stratified into single-neuron (microlevel), neural population (mesolevel), and whole-brain (macrolevel) dynamics. Conventionally, it is usually assumed that smaller units on the lower scale (e.g., neurons or neural populations) coordinate, synchronise, and integrate their activity to generate higher-order patterns that shape global brain dynamics [7, 8].

Efforts to understand how these functional interactions relate to consciousness have frequently focused on pairwise couplings, using network-theoretic approaches to examine meso-to-macro transitions and their relevance to function and consciousness [9–16]. Beyond pairwise interactions, hypergraphs and simplicial complexes capture higher-order relationships that cannot be reduced to simple dyads [17, 18]. Relatedly, techniques such as principal component analysis (PCA), brain eigenmodes [19, 20], and oscillatory harmonic modes [21, 22] have been used to describe large-scale neural dynamics in terms of spatially extended, low-dimensional patterns, offering alternative frameworks to capture mesolevel integration beyond dyadic interactions. While these approaches often presuppose that higher-order, macroscopic dynamics arise from lower-scale interactions, implicitly invoking the notion of emergence, they typically do not set out to measure emergence itself. Moreover, assuming predefined, discretely stratified hierarchical scales, they fail to accommodate a *heterarchical* view of the brain’s dynamical organisation, where transient and overlapping interactions are not strictly contained within any single scale or pair of scales. Addressing this gap, the current study aimed to detect macroscopically organised neural dynamics, as well as quantify their degree of emergence.

A growing body of theoretical [23, 24], methodological [25–28], and empirical [29–35] work supports the notion that emergent higher-order interactions govern whole-brain functions and influence arousal and conscious states. Information-theoretic measures capturing synergy—a dimension of emergence that quantifies the information about a target variable that is present when considering multiple source variables jointly, and not in any source individually—have revealed that human brains exhibit higher synergy than non-human primate brains [36], and synergy is reduced in patients with disorders of consciousness [32]. Another study found that synergistic emergence is greater in human wakefulness than in states associated with diminished consciousness [37]. While these measures identify the degree of synergy present in higher-order interactions, they do not directly identify the emergent dynamical organisation of these higher-order interactions across spatial scales.

Here, we apply a new approach that defines emergence through *dynamical closure*, whereby the degree of emergence is quantified by assessing how statistically independent macroscopic processes are from their microlevel base(s).

To operationalise this, we employ an information-theoretic dimensionality reduction technique called Dynamical Independence (DI) [28]. DI identifies low-dimensional subspaces, termed *n*-macros (with *n* indicating the spatial scale), and quantifies emergence by assessing their dynamical closure. Formally, we measure the *dynamical dependence* (DD), which quantifies the extent to which future macro-states can be predicted from past micro-states. More precisely, an *n*-macro is deemed emergent to the degree that knowledge of the micro-states does not significantly improve the predictability of the macro’s future states beyond the degree to which the macro states already self-predict. Minimising DD across scales then identifies the *n*-macros that maximise dynamical *in*dependence, and thus, are maximally emergent at that scale. DI therefore captures a distinctive aspect of emergence separate from synergy: emergence through the dynamical *closure* of the macroscopic dynamics. We further extend the framework of DI by endowing it with a principled method to not only quantify emergent dynamics but also identify the structure of the higher-order landscape as it emerges, allowing us to transcend predefined hierarchical levels by computing emergence across all candidate scales in a system and capture its organisation [38].

Previously, using biophysically-plausible neural models, DI was shown to capture the modulation of emergent dynamical structure by incrementally varying functional integration and segregation [38]. To translate this approach to real-world data, here we determine the degree of emergence and the corresponding dynamical structure, as quantified by DI, in different states of consciousness; in particular wakefulness and anaesthesia. Anaesthesia-induced behavioural unresponsiveness is often used as a proxy for the loss or substantial alteration of consciousness [39]. Different anaesthetic agents produce distinct experiential effects: propofol and xenon typically abolish any reportable conscious experience, whereas ketamine often induces a dissociative, dream-like state [40, 41].

These differences in conscious states have been previously linked to changes in the level of cortical and thalamic activity [42, 43], or to changes in terms of oscillatory activity estimated with the power-spectral density (PSD) [44–46]. Propofol is associated with a strong increase in delta power also observed in sleep [47] and an increase in frontal alpha [48–50]. Xenon anaesthesia shifts the distribution towards delta frequencies without alpha [50]. In propofol and xenon, evidence suggests that the increase in delta power reflects the occurrence of bistable dynamics marked by alterations between neuronally active (‘up’) and neuronally inactive (‘down’) states [51–54]. By contrast, ketamine generally leads to a reduction in prominent oscillatory peaks, with alpha power dispersing into theta and high beta bands [44, 48]. Beyond these spectral shifts, anaesthetic-induced unconsciousness involves altered effective connectivity [55, 56], changes in signal diversity [57, 58], variations in the spectral exponent of the PSD [59], and shifts in measures related to criticality [60, 61]. While these established markers are useful indices of conscious level, they fall short of explaining how these anaesthetics impact emergent dynamics and their organisational structure unfolded across spatial scales. Developing new tools for the quantification and description of emergent dynamical structure is a key step toward understanding the relationship between emergence and consciousness.

Anaesthetic states thus provide a valuable testing ground for our approach. We categorise propofol and xenon conditions as ‘no conscious-report’ states, since subjects report no conscious experiences upon regaining responsiveness, whereas ketamine states were considered as ‘affirmed conscious-report’, based on the dream-like reports provided (post-hoc) by participants. By applying DI to open-source EEG data recorded during these conditions [45], we examined how emergent dynamics are impacted by a modulation or alteration of consciousness. In particular, we explore whether DI can sensitively distinguish not only between wakefulness and unconscious states (propofol, xenon), but also between wakefulness and altered conscious states such as those induced by ketamine.

## 2 Results

We analysed spontaneous EEG recordings from 14 healthy adults (7 females, aged 18–28), each of whom underwent a single anaesthetic protocol involving either propofol (*n* = 4), xenon (*n* = 5), or ketamine (*n* = 5). Prior to anaesthesia, all participants completed a baseline wakefulness recording session, lasting 10 minutes, which served as the comparator condition for subsequent analyses. These wakeful recordings were collected immediately prior to the administration of anaesthetic, within the same session, ensuring within-subject comparisons across conditions. Anaesthetic depth was titrated to unresponsiveness (Ramsay Scale 6), and EEG was continuously recorded throughout. Only the spontaneous EEG segments—before (wakefulness) and during unresponsive anaesthesia—were used in the present analysis. As previously reported [45], only participants in the ketamine group retrospectively affirmed any conscious experiences upon emergence; participants in the propofol and xenon groups reported no such experiences.

In the subsequent analysis we first quantified the *degree of emergence* by measuring the DD of optimised macroscopic variables (*n*-macros) on the underlying EEG sources across multiple spatial scales. Second, to characterise the *emergent dynamical structure*, we examined whether specific *n*-macros— low-dimensional macroscopic variables minimising dynamical dependence— consistently reappear across optimisation runs, indicating the presence of dominant, stable higher-order processes within each conscious state.

### 2.1 The Dynamical Dependence of Macroscopic Variables across Higher-order Scales

To measure the degree of emergence, we quantified the dynamical dependence (DD) of emergent *n*-macros on the underlying lower-scale process at multiple spatial scales. Here we define the time-series of the 46 source-reconstructed EEG sources as the lower scale which gives rise to emergent dynamics. In this formulation, DD can be interpreted as the extent to which information flows from the high-dimensional source-level system and continues to influence (in a predictive sense) the evolution of a given low-dimensional macroscopic process. Emergent *n*-macros are defined as the lower-dimensional subspaces of these sources that minimise DD (corresponding to maximising dynamical *in*dependence, DI), identified through repeated optimisations (*N* = 100 runs, see Section 5.5). Due to the non-convex nature of the optimisation landscape, different runs may converge on distinct (local) solutions, reflecting the presence of multiple near-optimal emergent *n*-macros at each scale (see also Section 5.5 for further discussion). This results in 100 *n*-macros per scale (2≤*n*≤9), per participant (*N* = 14). Thus, for each scale and across optimisation runs, we extracted distributions of DD values minimised by the optimisation procedure for the wake (W) and anaesthetic conditions (propofol, xenon, ketamine; P, X, K, respectively). In Fig 1 we present the comparison between the control condition (teal for wakefulness) and anaesthesia (orange for propofol, yellow for xenon, lilac for ketamine).

**Fig. 1.**
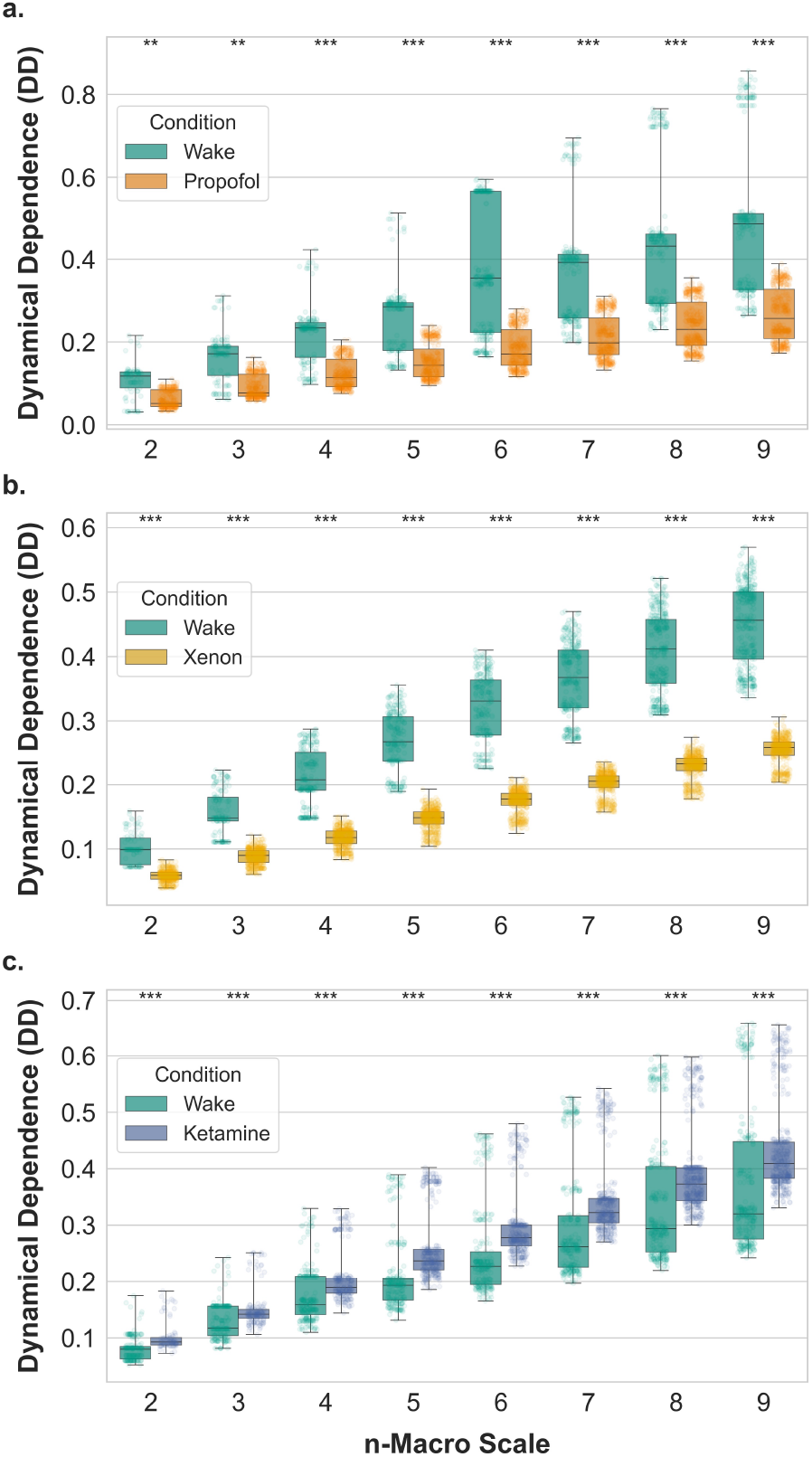
Dynamical Dependence across global states of consciousness and macro scales. The figure shows the distribution of Dynamical Dependence (DD) values across different macro-scales (2≤*n*≤9) for three anaesthetic conditions compared to wakefulness. *a*. Wake versus propofol, *b*. Wake versus xenon, and *c*. Wake versus ketamine. For each macro-scale, box plots display the median (central line), interquartile range (box), and full range (whiskers) of DD values. Individual data points are shown as semi-transparent dots, with teal representing wake measurements and condition-specific colours for each anaesthetic (orange for propofol, yellow for xenon, and lilac for ketamine). Statistical significance between wake and anaesthetic conditions is indicated by asterisks above each macro-scale (**: *p <* 0.001, ***: *p <* 0.0001). Higher DD values indicate stronger dynamical coupling—lower emergence— between higher-order *n*-macros and the lower-level sources, whilst lower values suggest increased emergence. The consistent differences across macro-scales demonstrate robust changes in the degree of emergence of macroscopic brain dynamics under different states of consciousness.

To determine whether the degree of emergence—quantified via dynamical dependence (DD)—differed significantly across conditions and spatial scales, we applied non-parametric comparisons of DD distributions at the individual participant level. These participant-specific comparisons were then combined across groups using a Z-score aggregation procedure (see Methods Section 5.6), which enabled us to assess whether DD values under anaesthesia were greater or smaller (in the sense of stochastic dominance [62]) than in the corresponding wake condition. This approach allows us to directly test for condition-related shifts in DD while accounting for inter-subject variability, and is statistically equivalent to Stouffer’s method [63] for combining independent z-values. Group-level significance was evaluated using false discovery rate (FDR) correction across macro scales.

The results illustrate a clear modulation of emergence across different anaesthetic agents, here measured via how emergent interactions in EEG are tied to lower-scale (source-level) processes. As shown in Fig 1a, compared to wakefulness, propofol yields significantly lower DD values across all examined spatial scales, indicating a more emergent state in propofol. This means that, under propofol, emergent interactions are relatively more independent of source-level processes.

Similarly, compared to wakefulness, xenon demonstrates a similar pattern to propofol, with significantly reduced DD values compared to wakefulness at all higher-order scales (Fig 1b), again pointing to increased emergence in xenon-induced anaesthetised state. In contrast, compared to wakefulness, ketamine shows the opposite effect where DD values across all *n*-macros are significantly higher (Fig 1c). This suggests that under ketamine, higher-order interactions are more closely tied to their underlying source-level processes, and are therefore less emergent.

### 2.2 Revealing Dynamical Structure of *n*-Macros across Conditions

To characterise emergent dynamical organisation across higher-order spatial scales, we examined whether optimisation runs within each participant tended to converge on similar macroscopic solutions at a given scale, *n*. Specifically, we assessed the extent to which distinct optimisations yielded convergent *n*-macros—measured via the subspace angle between each pair of resulting solutions (see Methods Section 5.5). For instance, two optimisation runs might produce *n*-macros *M*_1_ and *M*_2_, respectively. These *n*-macros could be identical (near-zero similarity) or exhibit a degree of dissimilarity. We quantified this similarity by measuring the subspace angle between *M*_1_ and *M*_2_. Normalised subspace angles approaching zero indicate near-identical solutions, while larger angles reflect distinctly different solutions. We interpret a proliferation of distinct solutions as reflecting a *fragmentation* of the global emergence structure.

The resulting pairwise similarity matrices, derived from 100 optimisation runs per subject—and ordered by ascending DD—revealed that under wakefulness, many runs converged on identical or near-identical *n*-macros, forming clear blocks of high similarity. This was most evident at lower-to-intermediate spatial scales but persisted, albeit with some dispersion, at higher spatial scales. In contrast, under propofol (Fig 2), this convergence deteriorated markedly. The similarity matrices showed greater dispersion, reflecting increased heterogeneity and *fragmentation* in the emergent solutions; this is indicative of flatter, less stable optimisation landscapes without dominant basins of attraction (locally minimal-DD *n*-macros).

**Fig. 2.**
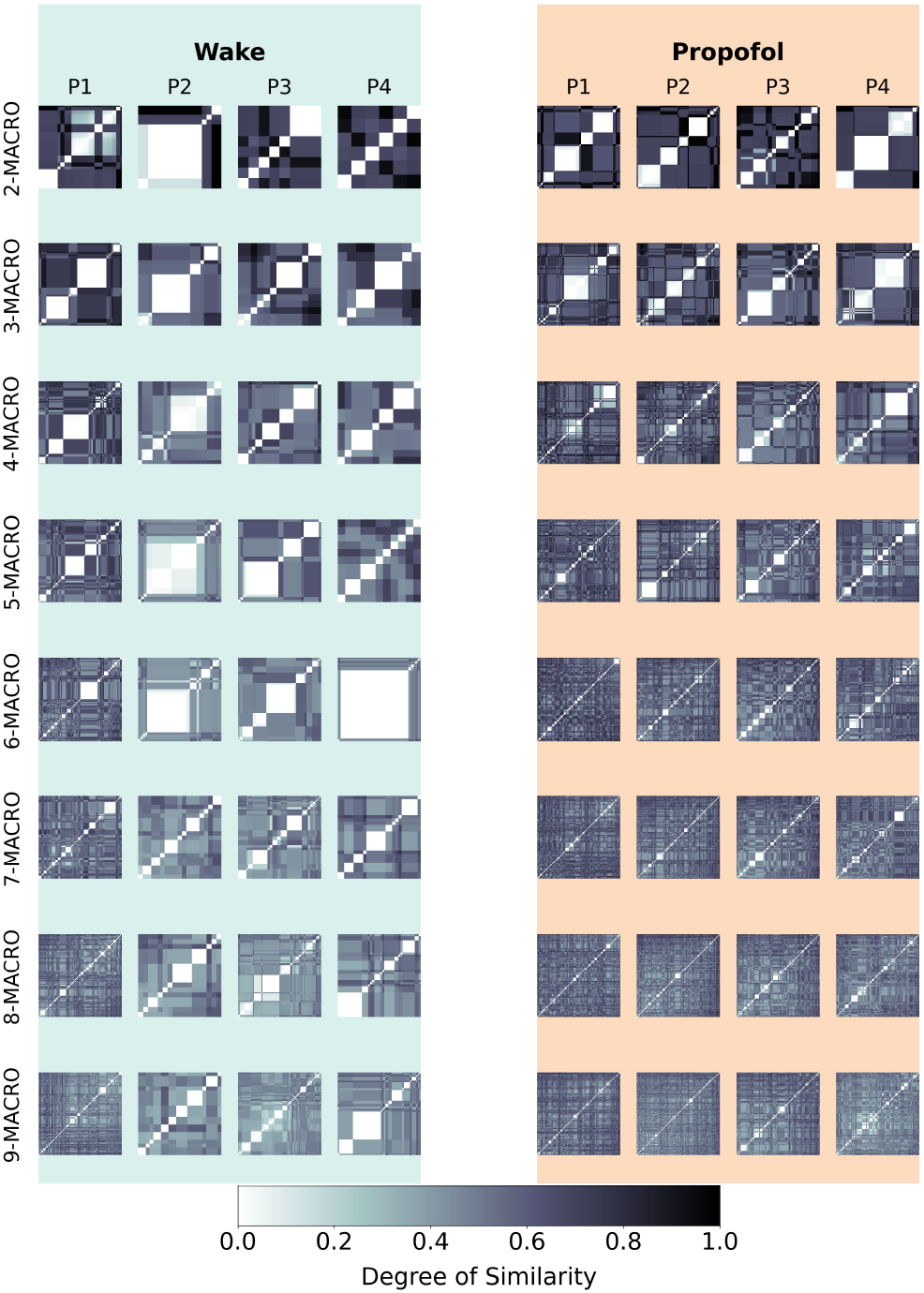
Propofol-induced Changes in Emergent Dynamical Structure on Higher-order Spatial Scales. Similarity matrices displaying the pairwise distances (subspace angles) between *n*-macros across optimisation during Wake (left) and propofol (right) conditions. The colour scale represents the principal angles (see Methods Section 5.5) between *n*-macros where greater dissimilarity between *n*-macros *M*_1_ and *M*_2_ is indicated with darker colours (black = orthogonal), and lighter colours signify greater similarity (white = identical). Both the *x*-axis and *y*-axis of the matrix correspond to the number of optimisation runs.

Importantly, we should not interpret this fragmentation in terms of the system selecting between competing *n*-macros at a given spatial scale, but rather that multiple emergent *n*-macros co-exist *simultaneously*, none of which necessarily dominate the dynamical organisation. This proliferation of distinct, co-existing emergent processes offers a precise characterisation of the fragmentation of emergent dynamics under anaesthesia.

Comparable findings were confirmed in the case of xenon, another anaesthetic condition associated with unresponsiveness and lack of reportable conscious experience. Compared to wakefulness, xenon produced marked fragmentation in emergent structure across all scales (Fig 3, with increasing divergence at higher macroscales. In contrast, under ketamine (Fig 4), the emergent structure more closely resembled that of the wake condition. Blocks of convergence persisted across most higher-order spatial scales, suggesting relatively preserved emergent dynamical organisation. Given that ketamine supports altered conscious experience rather than complete suppression, this finding supports preserved reports using the same dataset of experience in prior reports during dissociative states [45].

**Fig. 3.**
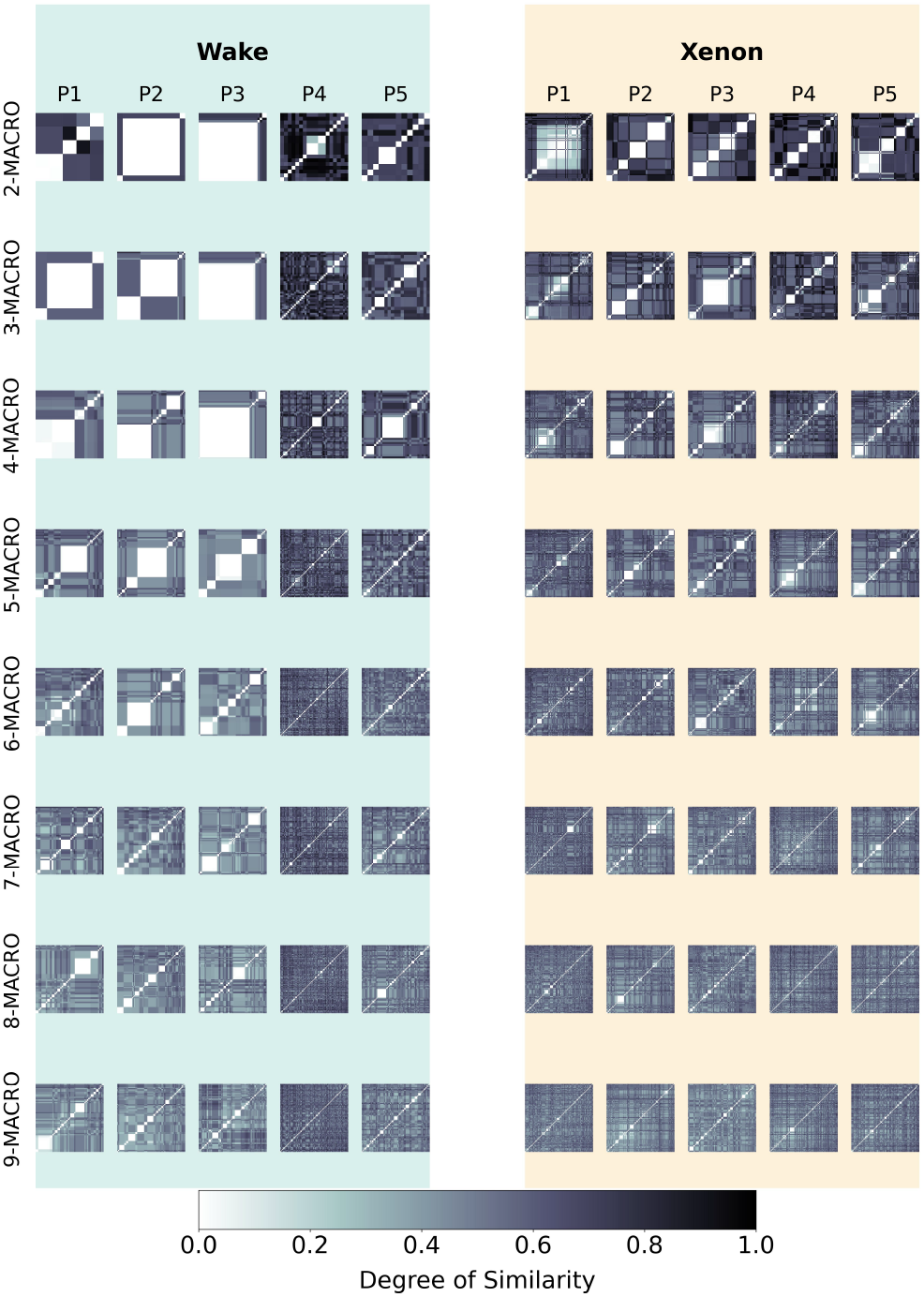
Xenon-induced Changes in Emergent Dynamical Structure on Higher-order Spatial Scales. Similarity matrices displaying the pairwise distances (subspace angles) between *n*-macros across optimisation during wake (left) and xenon (right) conditions. The colour scale represents the principal angles (see Methods Section 5.5) between *n*-macros where greater dissimilarity between *n*-macros *M*_1_ and *M*_2_ is indicated with darker colours (black = orthogonal), and lighter colours signify greater similarity (white = identical). Both the *x*-axis and *y*-axis of the matrix correspond to the number of optimisation runs.

**Fig. 4.**
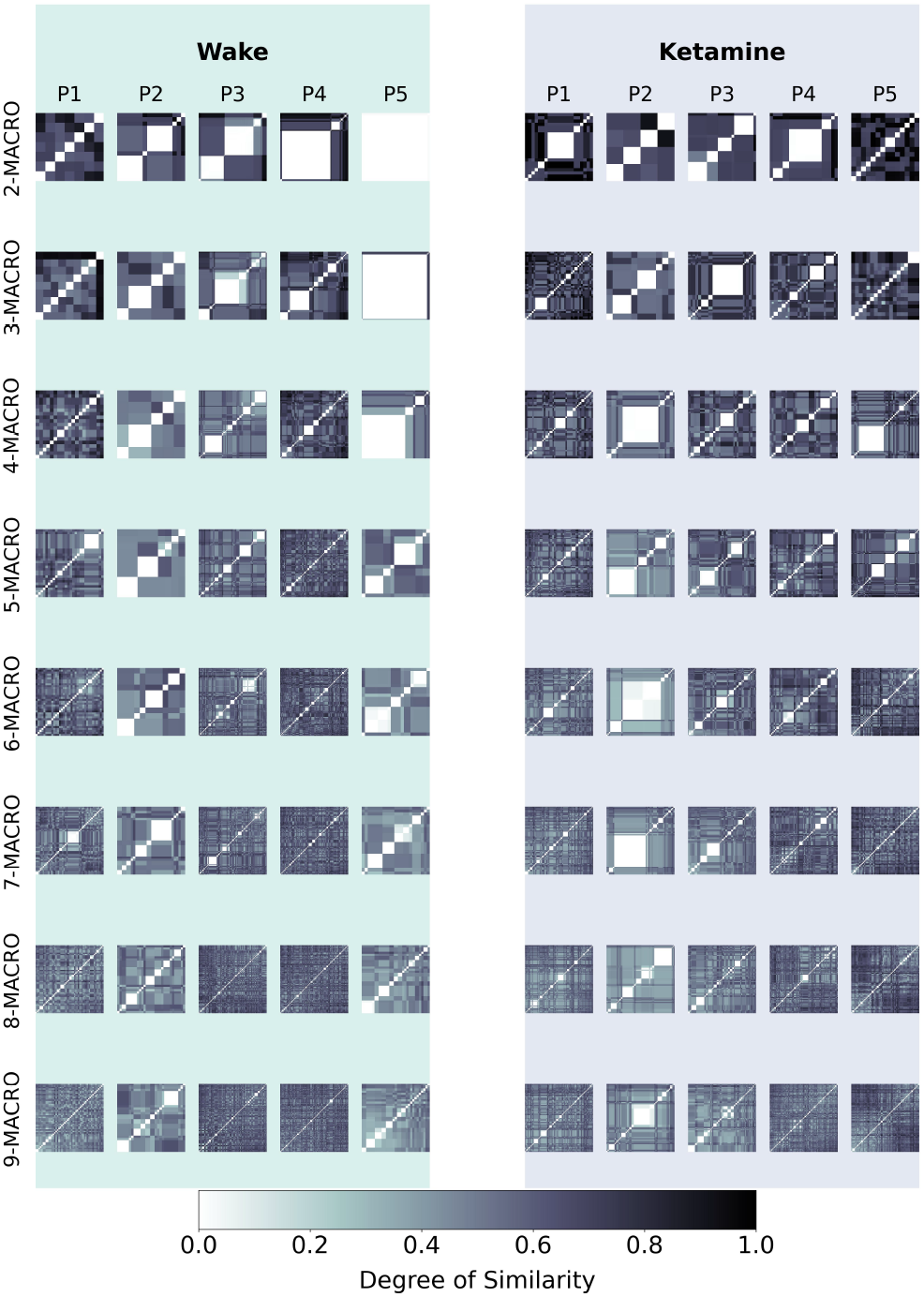
Ketamine-induced Changes in Emergent Dynamical Structure on Higher-order Spatial Scales. Similarity matrices displaying the pairwise distances (subspace angles) between *n*-macros across optimisation during wake (left) and ketamine (right) conditions. The colour scale represents the principal angles (see Methods Section 5.5) between *n*-macros where greater dissimilarity between *n*-macros *M*_1_ and *M*_2_ is indicated with darker colours (black = orthogonal), and lighter colours signify greater similarity (white = identical). Both the *x*-axis and *y*-axis of the matrix correspond to the number of optimisation runs

**Fig. 5.**
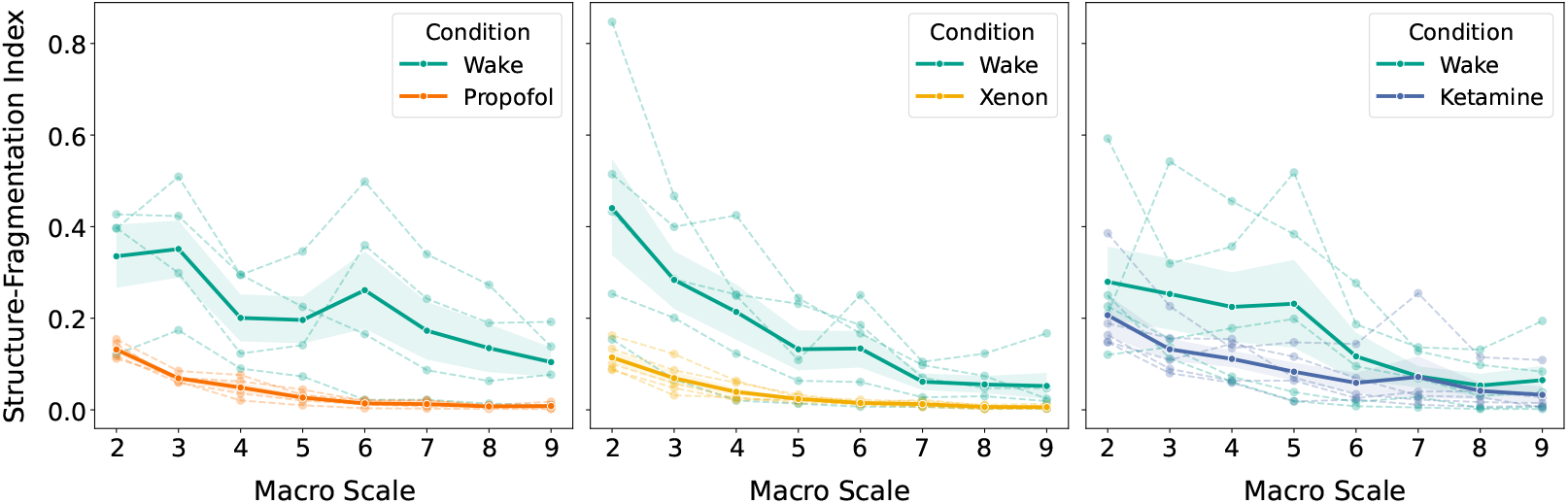
Emergent dynamical structure across wake and anaesthetic conditions (propofol, xenon, ketamine) Each panel displays the proportion of near-zero similarity values (*<* 0.0001) across 100 optimisations at each macroscale (2≤*n*≤9) for individual subjects under wakefulness (teal) and one anaesthetic condition. We consider this the *Structure-Fragmentation (SF) Index* of the emergent dynamical landscape. Dashed lines connect points from the same subject, allowing within-subject comparisons across conditions and scales. Group-level means are overlaid with 90% confidence intervals. Across all panels, wakefulness consistently exhibits a higher SF index (proportions of near-zero similarity values), indicating greater structure in emergent macroscopic landscape. Under propofol (left), a marked reduction in SF index is observed, particularly at intermediate-to-coarse scales, suggesting widespread fragmentation and instability in the emergent dynamical landscape. Xenon (middle) similarly shows degraded structure relative to wake, with pronounced divergence at higher-order scales (*n*≥7), reflecting scale-selective disruption and fragmentation of emergent organisation. In contrast, under ketamine (right), the structure of the emergent landscape macroscopic is relatively preserved, with trajectories closely tracking the wake condition across all scales. Although no macro scales survived FDR correction in post-hoc comparisons, the consistent directional trend of wake *>* anaesthesia is compatible with the hypothesis that emergent structure is more stable and organised in conditions permitting conscious experience.

To quantify these patterns, we introduce a *Structure-Fragmentation (SF) index* that computes the proportion of near-zero similarity values (*<* 0.0001) in 100×100 matrices across the eight coarse-grained macro levels. Using repeated-measures ANOVA with *subject* as a random factor, we tested for main effects of macroscale, condition (Wake vs Anaesthesia), and their interaction, followed by FDR-corrected post-hoc Wilcoxon tests to determine the directionality of the condition effect.

Compared to wakefulness, across all three anaesthetic agents—ketamine, propofol, and xenon—we observed highly significant main effects of macroscopic spatial scale on the SF index. Emergent structure varied substantially across scales in all cases: propofol (*F* (7, 21) = 9.99, *p <* .0001, *η*^2^ = 0.769), xenon (*F* (7, 28) = 16.38, *p <* .0001, *η*^2^ = 0.804), and ketamine (*F* (7, 28) = 15.35, *p <* .0001, *η*^2^ = 0.793). These findings robustly confirm the scale-sensitivity of structure, independent of condition.

The main effect of condition (Wake vs Anaesthesia) was significant for propofol (*F* (1, 3) = 10.65, *p* = .0470, *η*^2^ = 0.780), significant for xenon (*F* (1, 4) = 9.16, *p* = .0389, *η*^2^ = 0.696), and not significant for ketamine (*F* (1, 4) = 2.52, *p* = .188, *η*^2^ = 0.386). These findings suggest that propofol and xenon alter the overall structure and organisation of emergent dynamical landscape, reducing the proportion of near-zero similarities between *n*-macros compared to the wakeful state, whereas ketamine does not produce a reliable global difference suggesting preserved global organisation during dissociative states.

Most notably, the macro × condition interaction was again significant for xenon (*F* (7, 28) = 5.65, *p* = .0004, *η*^2^ = 0.585), with a trend for propofol (*F* (7, 21) = 2.42, *p* = .056, *η*^2^ = 0.446), and non-significant for ketamine (*F* (7, 28) = 1.21, *p* = .332, *η*^2^ = 0.232). These results indicate that xenon, and to a lesser extent propofol, modulate emergent structure in a scale-selective manner. In these regimes, the SF index is substantially higher during wakefulness than anaesthesia, particularly under xenon and propofol. This supports the hypothesis that wakeful brain states are characterised by greater macroscopic structure in emergent dynamical landscape; especially at coarse levels of description.

Together, these findings highlight the robustness of scale-dependence in emergent structure and suggest that scale-selective fragmentation of the structure of the emergent landscape is a key feature of the anaesthetic state, particularly under xenon. The pattern observed at higher macroscales under ketamine points to a potentially preserved or selectively altered macroscopic dynamical organisation, distinguishing it from GABAergic and NMDA-antagonist anaesthetics.

### 2.3 Source Contributions to Emergent *n*-macros

As an exploratory analysis, we next computed the contributions of individual sources to the *n*-macros to better understand the relationship between the higher-order interactions captured by *n*-macros and the lower-scale sources from which they emerge. Figs 6,7, and 8 illustrates the anatomical source-contributions to emergent *n*-macros above what would be expected by chance (please refer to Methods Section 5.5 for details on how we computed the source contributions and Methods Section 5.8 for statistical analysis).

**Fig. 6.**
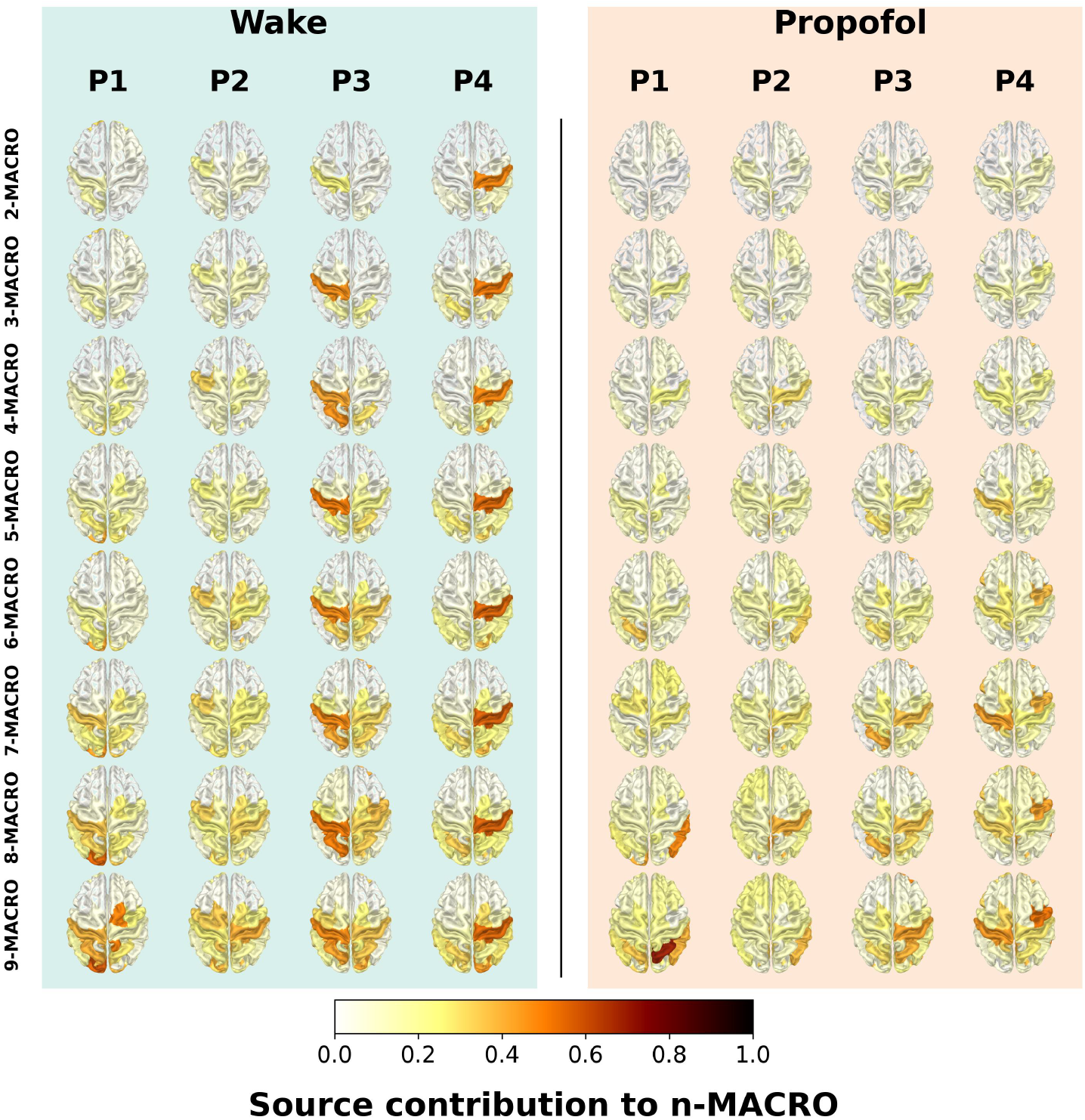
Source Contribution to Higher-order Interactions in Propofol v Wake. Single-region contributions per macro across subjects during propofol-induced anaesthesia and corresponding wake states. Each row represents an *n*-macro, and each column represents an individual subject. Colour intensity reflects the beta statistic (within the range [0, 1]), indicating the strength of each source’s contribution to the *n*-macro above chance. Zero equates to contributions expected by chance. The higher the values the stronger the contribution (refer to Methods Section 5.5 for details on beta statistic computation).

**Fig. 7.**
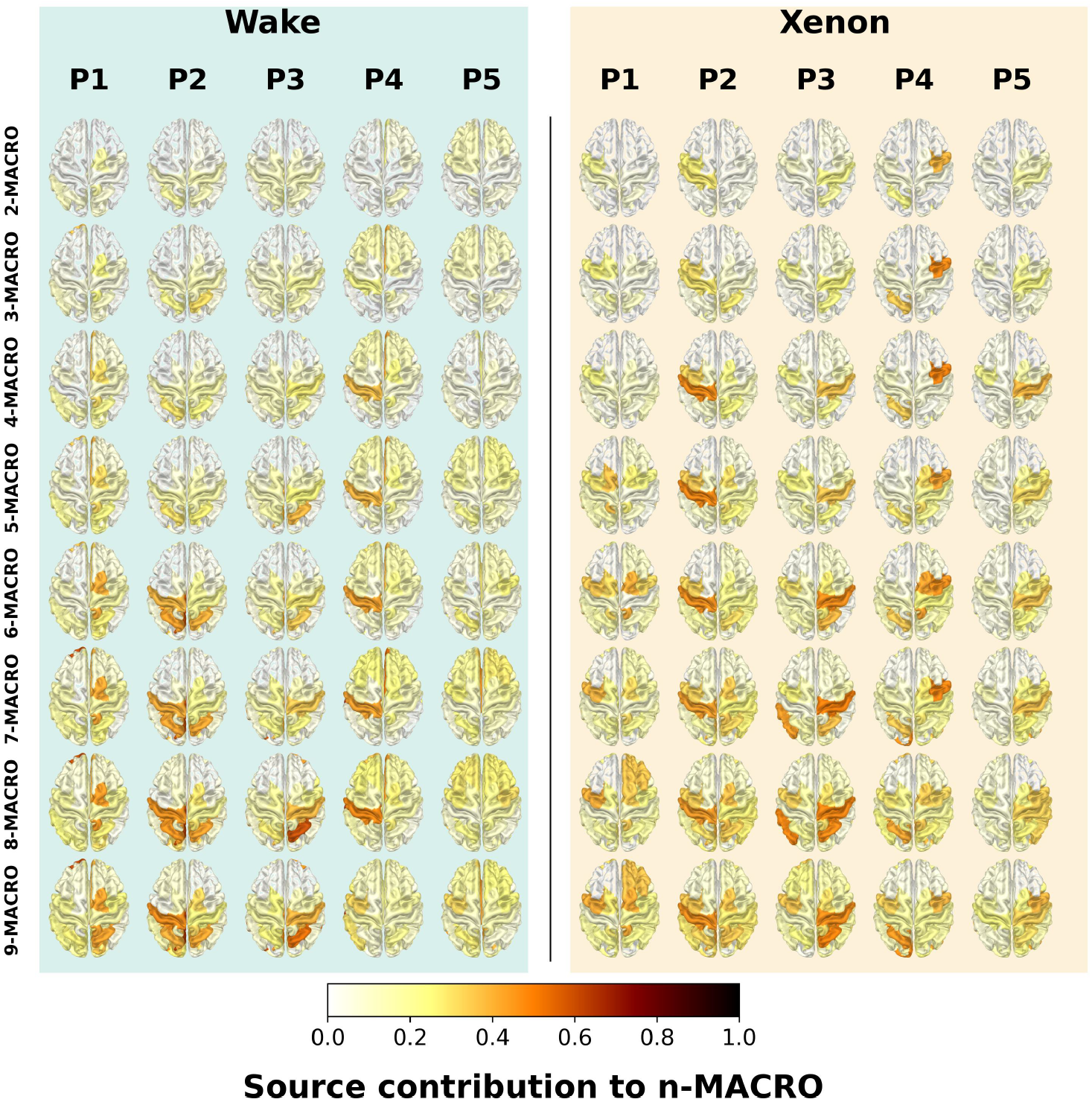
Source Contributions of Higher-order Interactions in Xenon v Wake. Single-region contributions per macro across subjects during xenon-induced anaesthesia and corresponding wake states. Each row represents an *n*-macro, and each column represents an individual subject. Colour intensity represents the Beta statistic (within the range [0, 1]), indicating the strength of each source’s contribution to the *n*-macro above chance. Zero equates to contributions expected by chance. The higher the values, the stronger the contribution.

**Fig. 8.**
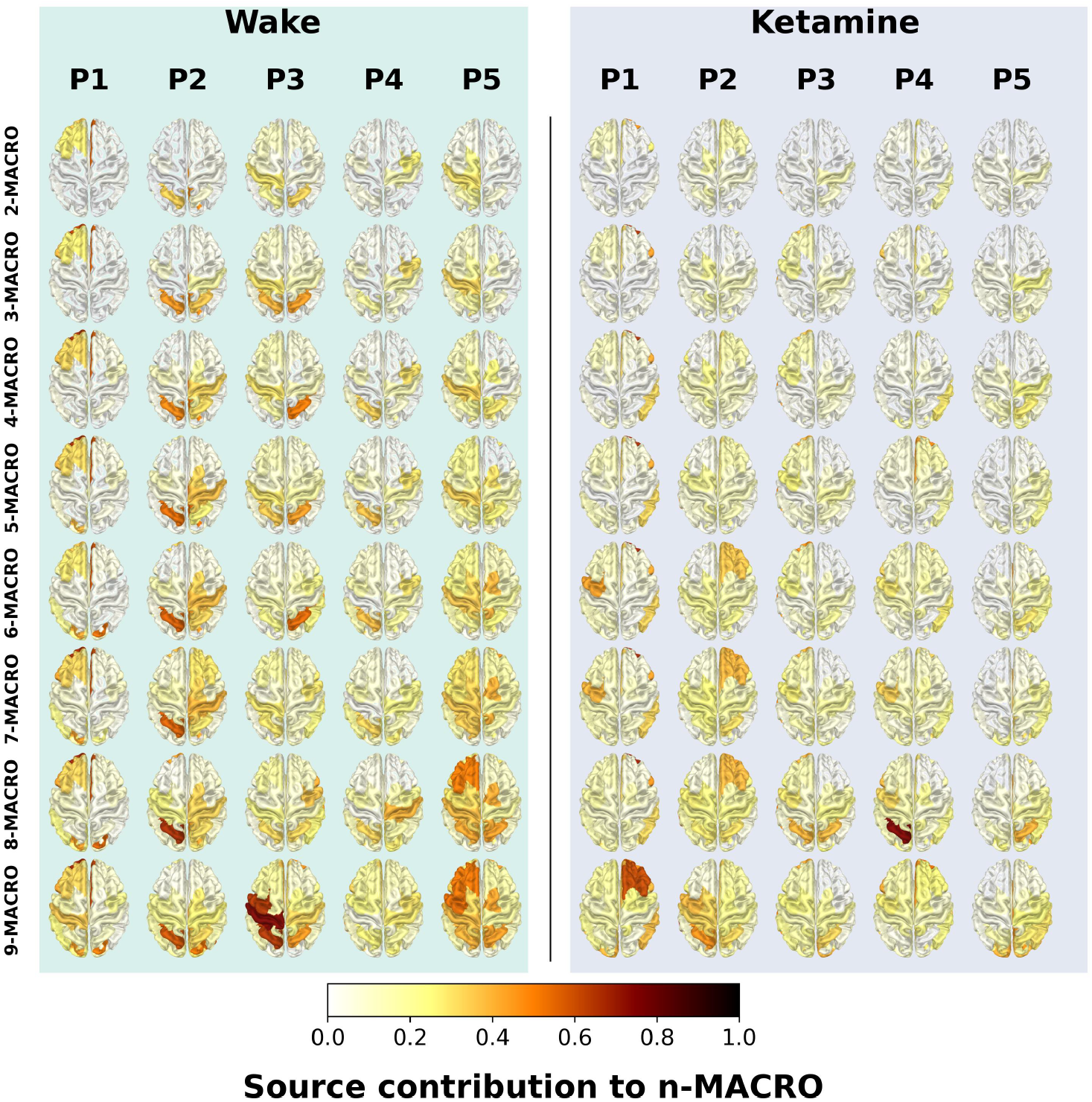
Source Contribution of Higher-order Interactions in ketamine v Wake. Single-region contributions per macro across subjects during ketamine-induced anaesthesia and corresponding wake states. Each row represents an *n*-macro, and each column represents an individual subject. Colour intensity reflects the beta statistic (within the range [0, 1]), indicating the strength of each source’s contribution to the *n*-macro above chance. Zero equates to contributions expected by chance. The higher the values, the stronger the contribution.

Wakeful participants consistently exhibit well-defined sets of sources of high contribution that persist or intensify at higher *n*-macro scales, reflecting a spatially-localised contribution of specific sources to emergent *n*-macros (Fig 6). Notably, participants P2, P3, and P4 display distinct localised patterns of strong contributions that overlap with the parietal cortex and extending across multiple scales. In two participants—P1 and P2—this contribution from the posterior and parietal regions is increased in higher scales. In contrast, the same participants under propofol show that these spatially concentrated patterns are less evident, yet a similar set of sources with high contribution over posterior regions is observed across higher spatial scales (*>* 6), which closely resembles wakefulness but is diminished in intensity. In general, the contributions appear more uniform with no clear prominence of specific sources. We note that in wakefulness, P3 and P4 exhibit strong lateralisation across the motor cortex, a pattern that becomes less pronounced and more bilateral with increased higher scales.

Compared to wakefulness, xenon does not exhibit stronger source contributions across higher-order scales. This marks a notable departure from the pattern observed under propofol, which does show increased higher-order contributions, suggesting a potential divergence in how these two anaesthetics affect large-scale brain dynamics. In both conditions, source contribution patterns vary across participants, but overall contributions tend to increase with scale and are similarly concentrated in the parietal cortex for both, wakefulness and xenon. However, in two out of five participants, frontal source contributions are reduced under xenon relative to wake. Some lateralisation is also evident across individuals in both states. These results suggest that while xenon preserves aspects of the posterior dynamics observed during wakefulness, this activity may be weaker, with reduced frontal contributions helping to distinguish the two states. The broader implications of these findings are discussed in the Discussion section.

Lastly, compared to wakefulness, ketamine shows a different pattern of source contributions across higher-order scales. While wake continues to exhibit increasing source contributions with scale—typically concentrated over posterior regions (with the exception of participant P1)—this pattern is less pronounced under ketamine. Specifically, ketamine displays lower overall increases in source contributions across scales and greater spatial heterogeneity. Notably, three out of five participants (P1, P2, and P4) show increased contributions in frontal regions under ketamine, contrasting with the posterior focus seen in wake. These findings suggest that ketamine differs from wakefulness not only in the degree of emergence and emergent dynamical structure, but also in the spatial localisation of their underlying source contributions.

Before proceeding to group-level comparisons across conditions, we note that the individual-level statistical analysis revealed that several regions showed significant contributions to *n*-macros after correction for multiple comparisons. These significant patterns are visualised in Supplementary Figures 5–7, which provide participant-level cortical maps for each condition, and each macroscale. Thus, although the following group-wise comparisons face stricter correction burdens, clear significant and localised source-level effects are already evident at the single-scale, single-subject level.

To complement the single-subject analysis, we evaluated group-level differences using a similar statistical approach as with the DD values in Fig 1. Specifically, we applied the grouped Mann–Whitney U-test separately to each source *i*, allowing us to determine whether the source’s contribution (*β*-scores) to an *n*-macro differs significantly between wakefulness and anaesthesia. As before, the test first compares each participant’s wakeful data against the corresponding anaesthetic data and then aggregates—across participants in the same anaesthetic condition—the U statistics and Z-scores using the methods defined in subsection 5.8 of the Methods Section. Notice that now the tests are *not* independent across sources, and thus we employed an FDR-based multiple comparison correction that assumes dependence of hypotheses.

Figure 9 visualises the resulting effect sizes: red indicates sources contributing more strongly to *n*-macros during anaesthesia, while blue highlights sources with stronger contributions under wakefulness. Although these effect size differences suggest potential distinctions in how each anaesthetic condition alters the spatial profile of emergent dynamics, none of these effects survived correction for multiple comparisons and should therefore be interpreted with caution. Nevertheless, the significant individual-level patterns (Supplementary Figures 5–7) provide complementary evidence of source-specific reorganisation across states.

**Fig. 9.**
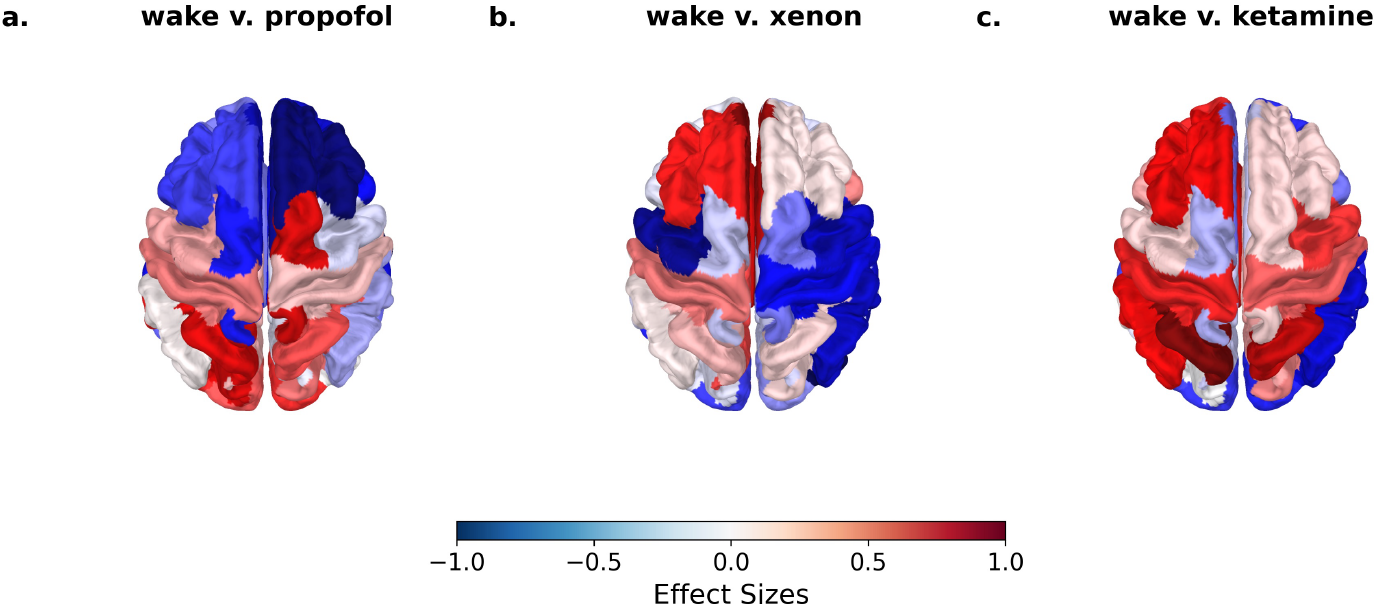
Contrasts between source contributions across all *n*-macros and conditions. Source contributions to *n*-macros across all higher-order scales (2≤*n*≤9) in the wake condition compared to (a) propofol, (b) xenon, and (c) ketamine. Red denotes sources where the wake condition shows greater contributions than the anaesthetic condition, while blue indicates greater contributions in the anaesthetic condition. A grouped Mann-Whitney U-test with FDR (with dependence) correction was used. After grouped-level corrections, no significant effects across conditions survived. This test assesses the stochastic dominance of the wake condition over the corresponding anaesthetic condition.

Compared to wakefulness, all anaesthetic conditions show alterations in the spatial distribution of source contributions to higher-order dynamics. For propofol, participants during wakefulness show that contributions are concentrated around posterior regions (Fig 9a), consistent with individual-level patterns. Under propofol, these contributions appear to shift toward frontal regions, as suggested by negative effect size values. In the xenon condition, posterior contributions during wake are not as prominent, and xenon introduces greater lateralisation and additional contributions from frontal and parietal sources (Fig 9b). For ketamine, source contributions also appear focused in posterior areas during wake, but with reduced frontal involvement under ketamine relative to wake (Fig 9c)—a pattern opposite to that observed under propofol and only weakly present in xenon. However, substantial between-subject variability, particularly evident even during wakefulness, highlights the need for larger sample sizes to more robustly characterise these patterns.

## 3 Discussion

In this study, we quantified the degree of emergence using dynamical independence and introduced the SF index to characterise the fragmentation of the emergent landscape. The SF index specifically captures the prominence of emergent n-macros across higher spatial scales. In parallel, we assessed the spatial embeddedness of these n-macros to further characterise the structure of emergent dynamics across wakefulness and three anaesthetic conditions.

Our findings indicate that across higher-order spatial scales, propofol and xenon were characterised by higher emergence compared to wakefulness, whilst ketamine was characterised by reduced emergence.

Given the common conflation of complexity and emergence in the neuroscience literature, it is understandable that one might intuitively expect indices of emergence to track with signal diversity [58] and algorithmic complexity [64] which are often heightened during wakefulness and reduced under anaesthesia [57, 58, 65]. However, we caution against this assumption; algorithmic complexity and emergence represent fundamentally different aspects of system organisation. Within the DI framework, emergence is operationalised through dynamical closure rather than complexity. This distinction is important for interpreting our results.

In the dynamical-closure framework, a process is deemed emergent once its future can no longer be improved by considering the past of its microscopic constituents. Crucially, this criterion speaks only to independence from the lower-level base, not to how well the macro predicts itself. An *n*-macro can therefore be perfectly emergent yet *internally* incoherent—indeed, it might look like near–white noise. What matters for functional relevance, however, is whether such macros also exhibit coherent *internal* dynamics. Although the present study focuses on an aspect of emergent dynamical structure by way of the emergent landscape revealed by recurring, prominent *n*-macros (Figs. 2–4)—we did not evaluate the intrinsic structure of each macro, leaving that question for future work.

This shift of focus from the degree of emergence to aspects of emergent dynamical structure motivates our two-part analytic strategy: we quantify (i) the degree of emergence (closure from the micro-level) and (ii) the organisation and structure of the emergent landscape across spatial scales. Treating both facets as complementary is essential for a full account of emergence in brain dynamics.

In light of this, it is most informative to view our findings through the lens of emergent *structure*. While propofol and xenon exhibit higher emergence—indexed by lower dynamical dependence—they lack prominent *n*-macros across scales, implying fragmented dynamical organisation. Conversely, wakefulness (and, to a lesser degree, ketamine) shows dominant, recurring *n*-macros that point to structured emergent landscape. We therefore developed SF index—quantifying the frequency with which an *n*-macro re-emerges across optimisation runs—as a practical proxy for robust higher-order structure.

Within the optimisation landscape, multiple *n*-macros can co-exist at any scale; fragmentation signals not competition among alternatives but the absence of a deep basin of attraction. Repeated recovery of the same *n*-macro indicates such a basin and thus dominant dynamics, whereas scattered minima point to a disorganised, noisy landscape. Similarity-matrix ‘white’ blocks visualise these basins, enabling our extended framework to separate structured emergent landscapes from high-dimensional noise, using a simplified SF index, and to pinpoint candidate neural correlates of conscious state.

Our findings challenge the intuitive relation between “more emergence” and conscious level. Propofol and xenon exhibit high dynamical closure but fragmented emergent landscapes, whereas wakefulness—and partially ketamine—show lower closure yet structured emergent dynamical landscapes. This suggests that the degree of emergence is only part of the story; macros must remain partially coupled to their microscopic dynamical substrate to be functionally relevant [66].

An implication of our findings is that conscious states eschew a clean and full *separation of scales* between micro and macro, which would be reflected in (near)-perfect dynamical closure/independence ((near)-zero DD). Strikingly, the *scale-integration* of the brain’s dynamics, at least under conscious conditions, contrasts strongly with engineered systems such as computers, where a strict hardware-software separation of scales is enforced by design. The general idea emerging here is that engineered systems enforce a neatly separated hierarchy of scales, making design easier, whereas evolved biological systems in certain states, like wakeful consciousness, operate in a scale-integrated manner across multiple levels of description[67–71].

The ketamine condition underscores this point. Despite reduced closure (emergence), ketamine preserves a higher SF index (moderate emergent structure) and is accompanied by vivid, dream-like reports [45]. This convergence between emergent structure and subjective experience further supports structure and scale integration—not level of emergence *per se*—as a predictive marker of consciousness. These results suggest a “sweet spot” of emergence—where macroscopic processes are sufficiently closed to form structured multiscale dynamical landscape yet are still coupled to their micro-level sources. Future work should extend these findings by jointly evaluating each macro’s degree of dynamical closure—as we do here—*and* its self-predictive power, in order to examine it’s functional relevance. Many other interesting questions arise hereabouts: for example, does scale-integration also peak at this putative “sweet spot”, and how do these dynamical signatures relate to other powerful perspectives on complex systems, such as criticality [72, 73].

Although group-level source localisation did not survive multiple comparison correction, individual maps (Supplementary Figs. 5–7) revealed suggestive condition-specific patterns. During wakefulness, dominant *n*-macros consistently involved a posterior “hot zone”—encompassing occipital cortex, precuneus, and posterior cingulate—previously linked to conscious content in sleep and dreaming studies [74]. Changes in slow wave activity within this region, which reflect increased neuronal synchrony alongside disrupted cortico-cortical connectivity [75, 76] and reduced deterministic responses [77], correlate with the presence or absence of subjective experience during sleep. This highlights the possible crucial role of parietal regions in generating conscious experiences [78]. Under propofol this posterior weighting was disrupted and frontal contributions increased, echoing the well-known frontal-alpha shift of this anaesthetic [48, 49].

Taken together, these preliminary data suggest that the anatomical localisation of structured emergent dynamics shifts with state. Specifically, posterior cortices may dominate in conscious conditions, whereas deep anaesthesia may recruit more anterior generators. Because the sample size was modest and inter-individual variability substantial, higher-resolution techniques (MEG, sEEG, iEEG) will be required to confirm whether posterior hot-zone engagement is a reliable signature of scale-integrated macros in conscious brains.

### 3.1 Comparison with alternative approaches to higher-order interactions

By going beyond measuring the quantity of emergence, we identified the underlying emergent dynamical structure in the electrophysiological data across global conscious states. Our method addresses the broader challenge of defining the multiscale functional organisation of biological brains directly from data. Building on early ideas [79] and recent developments in neuroscience [80] and biology [81], we emphasise that global brain states (particularly of consciousness) involve reorganisation of the multiscale functional architecture of brain dynamics.

Neural activity has previously been shown to organise into low-dimensional subspaces [82, 83]. Our study builds on this work to demonstrate that DI provides a principled way to capture both the degree of dynamical closure across multiple scales, and to identify emergent dimensionally-reduced macroscopic processes directly from neurophysiological data. Our findings support a heterarchical view of brain organisation, consistent with the idea that higherorder scales remain partially dependent on lower-level dynamics [68, 79]. Furthermore, our results align with recent work suggesting that dimensionality reduction reveal higher-order interactions [83, 84], and they illustrate how DI can offer a unified approach for quantifying the emergence of higher-order interaction and capturing the macroscopic organisation of neural activity.

Our approach invites comparison with other information-theoretic measures of emergence designed for neuroscientific data [26, 27]. For instance, synergy-based causal emergence focuses on the additional information provided by higher-order variables in predicting future states beyond what is carried uniquely by the parts [27, 85–88]. These methods have identified synergistic higher-order interactions in various contexts, including neurodegeneration [33], ageing [89], resting-state fMRI in humans and macaques [31, 34, 36, 90], neural spike dynamics [91], and mouse neocortical cultures [92]. Despite recent extensions, information decomposition methods still face practical constraints when extending beyond small higher-order sets (*n >* 3) [93], necessitating secondary graph-theoretic strategies to glean insights from higher-order interactions [31, 34]. In this sense, synergy can be viewed as a specific case of higher-order interactions [6, 37, 94, 95], potentially distinct from the notion of dynamical closure at the heart of our DI-based definition of emergence.

Further, unlike *cliques, motifs*, or *simplicial complexes*—which are topological or graph-based structures used to represent higher-order interactions [9, 18]—*n*-macros serve as lower-dimensional subspaces that capture relevant multivariate dynamics. Conceptually similar to principal components (see Methods Section 5.5), *n*-macros highlight how multiple variables (for instance, brain regions) collectively interact, potentially offering a fresh perspective on *communication subspaces* [80, 96].

This DI-based approach can complement synergy-based methodologies, graph-theoretic analyses [97, 98], and simplicial complex frameworks [17, 18, 99], by identifying macroscopic observables directly from data rather than relying on predefined macro variables. Furthermore, because DI scales reasonably well with both system dimension and by virtue of the reduced dimensionality of *n*-macros, this approach circumvents some of the in principle computational constraints inherent in standard information decomposition methods [37, 85] as mentioned above. Although recent work using *O-information* [34, 35, 100] seeks to overcome these constraints in synergistic interaction analysis, the direct discovery of *n*-macros via DI offers a distinct advantage when dealing with high-dimensional data.

### 3.2 Limitations and Future Directions

This study quantifies the structure of emergent dynamics by tracking (i) how strongly *n*-macros dominate higher-order scales (similarity matrices) and (ii) where those macros localise over brain sources. Because EEG suffers from volume-conduction artifacts, its spatial precision is limited even after source reconstruction; MEG, ECoG, or sEEG/iEEG will be better suited for fine-grained spatial localisation. Even so, our results show that EEG is sufficient to reveal scale-dependent emergent structure.

The modest sample size urges replication with larger cohorts, yet the cross-condition consistency of our similarity matrices (Figs 2–5) and the method’s single-subject resolution already indicate its potential it for clinical application, in contexts such as anaesthesia tracking or disorder-of-consciousness assessment, which could prove to be a useful tool for precision medicine. Gradient-descent optimisation proved robust, but complementary schemes (e.g. simulated annealing or genetic algorithms) will clarify whether fragmented emergent landscapes reflect physiology or search bias.

Future work should test whether the macros we detect align with Integrated Information Theory’s (IIT) posterior hot zone [101, 102] or Global Neuronal Workspace Theory’s (GNWT) [103, 104] fronto-parietal broadcast—ideally in adversarial, theory-driven designs [105, 106]. Augmenting DI with internal diagnostics—macro self-predictability, model-complexity indices (AIC/BIC), entropy rate, and persistence measures such as spectral radius or autocorrelation decay—will reveal whether candidate *n*-macros are genuinely self-determining, internally and externally structured processes or merely noise, thereby sharpening the empirical bridge between multiscale emergent structure and conscious experience.

## 4 Conclusion

Our study highlights a sweet-spot of brain organisation: consciousness thrives when emergent dynamics rise above local activity yet remain coupled across scales. Propofol and xenon push cortical activity toward highly decoupled macrodynamics that realise a more fragmented emergent dynamical landscape. Whereas wakefulness—and, to a lesser degree, ketamine–preserve a structured, scale-integrated emergent dynamical landscape despite lower emergence. By extracting these macrodynamics directly from neurophysiological, we uncover a heterarchical architecture in which partially scale-integrated emergent structures, not fully independent ones, sustain both the level and the richness of conscious experience. The DI framework thus offers a powerful, theory-agnostic lens to map the multiscale dynamical organisation onto cognitive conscious processing.

## 5 Methods

### 5.1 Data Overview

The present analysis solely utilised the spontaneous 61-channel EEG data of the open-source recordings described in [45]. Full details on participant selection, anaesthetic dosing, clinical monitoring, and retrospective assessments of subjective experience can be found in the supplementary material of that original work. Here, we offer a brief summary.

#### Participants

A total of 14 healthy adults (7 females), aged between 18 and 28 years, provided written informed consent and participated in the study, which was approved by the local ethics committee of the University of Liège, Belgium. All participants underwent medical and physical screening to exclude conditions that might interfere with anaesthetic administration. They were then assigned to one of three anaesthetic protocols: propofol (*n* = 4), xenon (*n* = 5), or ketamine (*n* = 5). Although the original dataset included transcranial magnetic stimulation (TMS) sessions, only the spontaneous EEG recordings were analysed in the present study [45].

#### Experimental Procedures

All testing took place at the Centre Hospitalier Universitaire (CHU) in Liège, where each participant received just one type of anaesthetic [45]. After administering a standard dose of anaesthetic—determined according to best practices for achieving unresponsiveness (Ramsay Scale score 6)—continuous spontaneous EEG was recorded. During pre-anaesthetic wakefulness, a 10-minute segment of spontaneous EEG was also obtained to serve as a baseline. Anaesthesia was then ceased, and participants were monitored until full responsiveness was regained.

Throughout the procedure, vital signs (e.g., blood pressure, oxygen saturation, exhaled CO_2_) and axillary skin temperature were tracked, and precautions (e.g., metoclopramide administration) were taken to minimise the risk of nausea. After participants had fully recovered, retrospective reports regarding any dream-like experiences or conscious thoughts during anaesthetic unresponsiveness were collected [45].

#### Conscious Reports

Participants who emerged from propofol and xenon anaesthesia reported no conscious experiences, with one xenon participant vaguely recalling a sensation upon emergence but without any specific memory. In contrast, subjects who emerged from ketamine-induced anaesthesia reported dream-like conscious experiences. These reports were collected one hour after emergence and categorised as either containing conscious experiences (‘affirmed conscious-report’ in our present study) or lacking such experiences (‘no conscious-report’ in our present study).

To provide an example, here is a quote from the supplementary information of the original publication [45]:

> “First I had the feeling of falling backwards into space. […] I then found myself in a futuristic spatial vessel like environment, discussing with two people I know about some abstract theoretical problem I am interested in, and we concluded that there was no solution, and that we disagreed on some fundamental points. At the end of this brief discussion I came up with the strong conclusion that everything I thought before was wrong, that nothing I believed was true anymore […]. At this stage, the space of my dream became slippery, I fell again with a feeling of shrinking space where everything became oblique, and I could see through the window a very large, white sphere in the intersideral space. […] I continued to fall and arrived in another room, still thinking that I had to find a solution to our theoretical problem using some kind of technology or way of calculation that did not exist yet; I had at that point a vision of a futuristic environment with a lot of machines, and some intense discussions with some unknown people about this mathematical problem. Then, reality started to fade and shrink further, the space of my dream flattened and virtually disappeared, and instead there was the apparition of something like a white and bright shape invading progressively the whole space. I would describe this scene as a beautiful end-of-the-world vision, with a real sensation of beauty, very impressive and strange, but no anxiety associated with it”

### 5.2 Preprocessing and Analysis of EEG Data

For the current study, the 61-channel spontaneous EEG recordings that were originally sampled at 1450 Hz were downsampled to 256 Hz for computational efficiency. Prior to downsampling, signals were high-pass filtered at 1 Hz to remove slow drifts, low-pass filtered at 250 Hz to attenuate high-frequency noise, and notch filtered at 50 Hz (and its harmonic at 100 Hz) to suppress line interference. After filtering, the data were re-referenced to the global average.

EEG preprocessing was followed by average referencing and projection of artifact-rejection vectors using MNE-Python. The continuous signals were segmented into 2-second non-overlapping epochs aligned with each labelled sleep or anaesthetic stage. This approach provided short, temporally precise trials suitable for source reconstruction and downstream multivariate analysis.

Epochs were treated as stationary multi-trial data under the assumption that each 2-second segment reflects a realisation from an underlying stationary stochastic process within a given conscious state. This assumption supports the application of Granger-causal inference [107] and is further justified by the stability of the fitted state-space models, ensuring that subsequent analyses are consistent and well-defined [108].

Epochs containing gross artifacts or bad channels were excluded upon visual inspection, resulting in a variable number of usable epochs across participants and anaesthetic conditions which are summarised in Table 1

**Table 1.** Participant-wise Distribution of Epoch Counts across Wakefulness and Anaesthetic Conditions.

| Wake | ketamine | propofol | xenon |
| --- | --- | --- | --- |
| 155 | 164 | 85 | 169 |
| 512 | 174 | 506 | 190 |
| 151 | 150 | 113 | 172 |
| 184 | 166 | 191 | 152 |
| 151 | 153 |  | 150 |
| 157 |  |  |  |
| 351 |  |  |  |
| 218 |  |  |  |
| 161 |  |  |  |
| 154 |  |  |  |
| 174 |  |  |  |
| 63 |  |  |  |
| 152 |  |  |  |
| 152 |  |  |  |

### 5.3 Source Reconstruction

For source reconstruction, a noise covariance matrix was computed from the cleaned continuous EEG using MNE’s empirical method. All source-level analyses were performed in a common template space using the `fsaverage` brain to ensure comparability across participants. A mid-resolution cortical mesh was generated using a fourth-order icosahedral subdivision (`ico4`), and a three-layer BEM model was constructed and solved using MNE-Python’s forward modelling routines. In the absence of individual MRIs, sensor locations were registered to the template head via the `fsaverage` transformation. Projecting to a shared source space reduces inter-individual anatomical variability and supports meaningful group-level analyses across conditions. A forward solution was then computed, with source-to-skull minimum distances constrained to 5 mm, resulting in a leadfield matrix mapping dipolar sources to sensor measurements.

To estimate cortical activity, a Linearly Constrained Minimum Variance (LCMV) beamformer was applied to each epoch using MNE-Python’s `make_lcmv` and `apply_lcmv_epochs` functions. Regularisation was set to 0.05 (5% regularisation and weight normalisation based on unit-noise gain), and a neural activity index (NAI) was used for depth compensation. The max-power orientation mode was selected to align each source with the dominant local field orientation. This yielded source-level time series for each epoch across the cortical mesh.

Subsequently, source estimates were parcellated into 46 anatomical regions using the reduced Human Connectome Project (HCP) atlas (Fig 10), accessed via MNE’s `read_labels_from_annot` function. Region-wise signals were extracted using `extract_label_time_course` in `mean_flip` mode, which averages within a label while preserving consistent polarity. Regions with zero-valued output (e.g., due to poor sensitivity or anatomical coverage) were excluded. The final data structure—epochs×time×regions—was stored per condition in both `.csv` and `.mat` formats for compatibility with subsequent dynamical independence analysis.

**Fig. 10.**
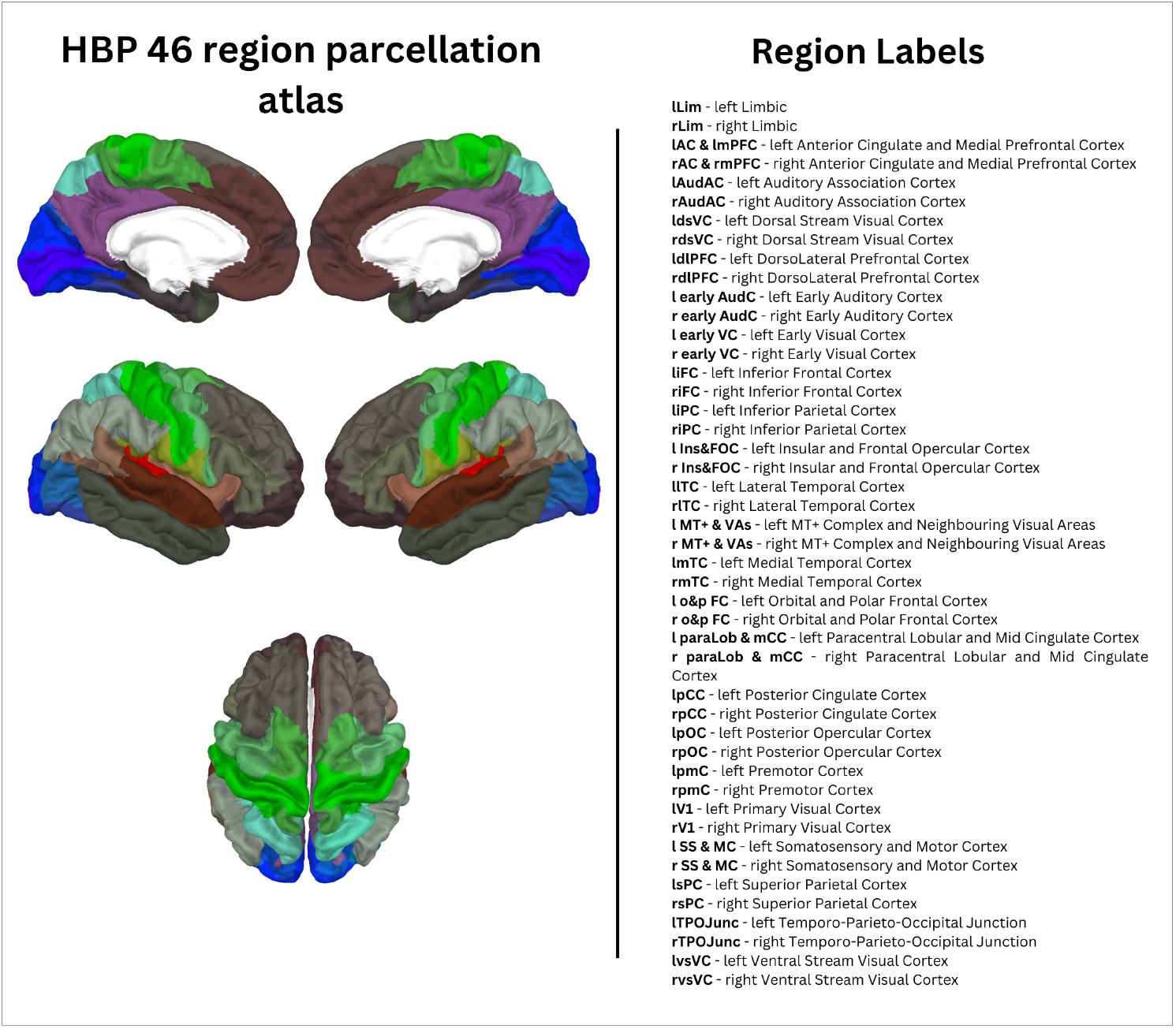
Reduced HCP Parcellation Atlas used for Source Reconstruction.

All preprocessing steps, including Z-score normalisation (i.e., mean subtraction and variance scaling), were performed separately for each participant and condition. The only exception was the Power Spectral Density (PSD) analysis (see Supplementary Information), which used raw (non-normalised) data to preserve absolute power differences. To ensure within-condition consistency, for all subsequent analyses wake EEG data were only averaged within each anaesthetic group and were not pooled across anaesthetic protocols.

### 5.4 Vector Autoregressive & State-space Modelling

To investigate emergent dynamical structure in the EEG data, Vector Autoregressive (VAR) models were fitted to the source-level signals using the MVGC2 toolbox [108, 109]. Model fitting was performed separately for each subject and condition. For each dataset, the optimal model order was selected by evaluating Akaike (AIC) and Bayesian (BIC) information criteria over a candidate range up to 30 lags. Given BIC’s tendency to underfit in practice, we selected AIC as our primary criterion, which typically resulted in model orders between 8 and 20.

VAR coefficients and noise covariances were then estimated via least-squares regression using MVGC2’s `tsdata_to_var` function, yielding a model of the form:

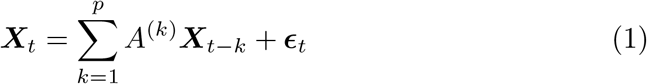

where ***X***_*t*_ is the *N*-dimensional multivariate source time series, *A*^(*k*)^ the autoregressive coefficient matrix at lag *k*, and ***ϵ***_*t*_ a zero-mean white noise (iid and serially uncorrelated) residual process with covariance matrix *V*.

To facilitate further analysis, each VAR model was converted into an equivalent state-space (SS) model in innovations form [110]. This was done using MVGC2’s `var_to_ss` routine, which constructs the companion-form state transition matrix *A*_*ss*_, observation matrix *C*, Kalman gain matrix *K*, and innovation noise covariance *V*_*ss*_. The system is described by the innovations state-space equations:

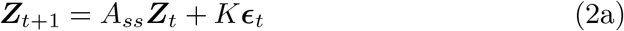

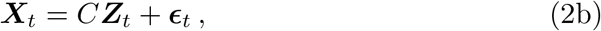

where ***Z***_*t*_ is the latent state vector of dimension *r*, which dynamically generates the observed activity **X**_*t*_ through linear mapping and innovation-driven temporal evolution.

Finally, spectral properties of the fitted systems were evaluated by transforming the VAR coefficients into a frequency-domain transfer function *H*(*ω*) using `var2trfun`, allowing for computationally-efficient calculation of dynamical dependence.

While not central to our main analysis, pairwise-conditional Granger-causal estimates were computed without statistical thresholding and provided in the Supporting Information.

### 5.5 Dynamical Independence: information-theoretic linear dimensionality reduction

After estimating the state-space parameters, we evaluated dynamical dependence (DD) to identify emergent macroscopic variables in the temporal domain.

In neuroscience, dimensionality reduction techniques are widely used to abstract key features of high-dimensional neurophysiological data—such as EEG recordings from electrodes or brain regions—by projecting them into lower-dimensional spaces that preserve structure of interest [111]. These methods often aim to capture global features like covariance, correlation, or temporal structure [112, 113]. Building on this idea, we frame Dynamical Independence (DI) as a linear, information-theoretic dimensionality-reduction technique, operationalised as an optimisation problem on a matrix manifold that seeks to minimise dynamical dependence [28]. Unlike Principal Component Analysis (PCA) [114, 115] or Maximum Autocorrelation Factors (MAF) [116, 117], DI is uniquely sensitive to the temporal dependencies in the data that underlie emergent structure.

A closely related idea from statistical physics is *coarse-graining*, which maps a high-dimensional microscopic system onto a lower-dimensional macroscopic description while retaining key system dynamics. In the context of neural systems, this approach enables us to extract dominant interaction patterns across spatial and temporal scales.

Here, we define a coarse-graining as a linear mapping *M* :ℝ^*N*^→ℝ^*n*^ where ***x***∈ℝ^*N*^ (e.g., 46-dimensional source-level data) is mapped to ***y*** = *M* ***x*** ∈ ℝ^*n*^, where ***x*** represents a variable in the full *N*-dimensional neurophysiogical phase space and ***y*** is a variable in the reduced-dimensional subspace. Intuitively, *M* can be viewed as a projection of the full neurophysiological phase space onto an *n*-dimensional subspace. To quantify DD efficiently, we use the following frequency-domain formulation:

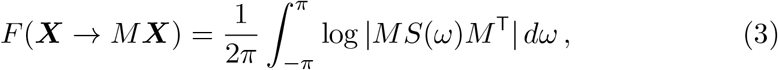

where ***X*** is the original high-dimensional source-level process, *M* the linear linear projection, and *S*(*ω*) is the *N* × *N* cross-power spectral density (CPSD) matrix at circular frequency *ω* ∈ [−*π, π*], which may be calculated from the VAR or SS model transfer function *H*(*ω*) and innovations covariance matrix *V* (to note, here we calculated the transfer function using both, the VAR and SS model parameters, where the analysis was performed on the SS model parameters. Calculating either VAR model parameters and converting them to SS model parameters, or calculating SS parameters directly, is theoretically equivalent for our analysis that relies on the transfer function *H*. The exhaustive sanity check employed here therefore also converged). This formulation captures how the low-dimensional process ***Y*** = *M* ***X*** inherits dynamical (or statistical) dependence from ***X*** across all frequencies (i.e., in the temporal domain).

In full generality, DD is invariant under non-singular transformations of both the microscopic and macroscopic spaces [28]. In the linear case addressed here, this means that we may identify the space of (linear) macrovariables with the *Grassmannian manifold* 𝒢_*N*_ (*n*) of *n*-dimensional linear subspaces of ℝ^*N*^ [118]. An implication of this is that we may restrict the linear mappings *M*, represented by *n*×*N* matrices where the row-vectors span the subspace in question, to be *orthonormal*; i.e., *MM* ^T^ = *I*. The space of such matrices is known as the *Stiefel manifold* 𝒱_*N*_ (*n*) [119]. We note that the mapping from orthonormal matrices to the corresponding Grassmannian manifold is many-to-one (orthogonal transformations Φ of ℝ^*n*^ taking *M* → Φ*M* leave invariant the linear subspace in ℝ^*N*^ spanned by the row-vectors of *M*), so that the “Stiefel parametrisation” of the space of macroscopic variables by *n*×*N* orthonormal matrices is redundant.

We now formally define an emergent *n*-macro:

#### Definition 1

(*n*-macro) An *n*-macro is an *n*-dimensional linear subspace of the *N*-dimensional source variable space. An *emergent n*-macro is an *n*-macro that locally minimises dynamical dependence^1^.

To compute emergent *n*-macros efficiently, we represent them as described above via the Stiefel parametrisation—i.e., as orthonormal *n* × *N* matrices *M* ∈ 𝒱_*N*_ (*n*)—and optimise the DD function (3) on this manifold. To improve computational tractability, we first minimise the proxy objective function:

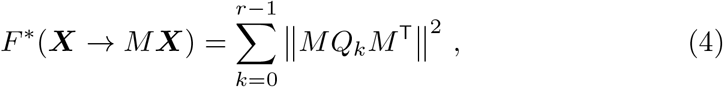

which stands as a computationally cheaper approximation to *F* (***X*** →*M* ***X***). Here *Q*_*k*_ = *CA*^*k*^*K*, where (*C, A, K*) are matrix parameters of a state-space model, and ∥ · ∥ denotes the Frobenius matrix norm. Crucially, the *Q*_*k*_ matrices are fixed for the given system, and may thus be pre-computed prior to optimisation of *F* ^*^(***X*** → *M* ***X***), which circumvents computing the state-space parameters at each optimisation step. This two-step optimisation enables efficient identification of emergent *n*-macros across multiple spatial scales [28, 38].

By optimising the objective function in (3), our method captures unique features in the data that differ from those identified by PCA or MAF. DI extends traditional variance-and dynamics-based methods in several key ways:

- **Scale Flexibility:**Like PCA, DI can identify low-dimensional subspaces at any scale 1≤*n*≤*N*. However, unlike PCA, which offers no notion or metric of emergence, DI explicitly quantifies emergence via dynamical dependence, providing a principled way to identify macroscopic variables that are not only lower-dimensional but also dynamically independent from their microscopic substrates.
- **Sensitivity to Dynamics:**Unlike PCA, which captures only contemporaneous variance, and MAF, which maximises 1-lag autocorrelation, DI explicitly incorporates the full temporal structure of the system, including lagged dependencies across multiple time steps. This allows it to identify subspaces that reflect the actual dynamics driving the evolution of the system.
- **Information-Theoretic Basis:**DI as presented here is implemented in terms of Granger causality, and as such inherits the information-theoretic interpretation of Granger causality [120, 121]; specifically DD may be interpreted as the information flow from the high-dimensional source-level system to the low-dimensional subsystem, offering a measure of (departure from) emergence of the dimensionally-reduced system.
- **Quantifies Source Contributions:**As we propose in Section *Characterising Source-level Contributions to n-macros*, DI admits a natural quantification of the contribution of individual sources to *n*-macros, facilitating analysis of spatial localisation of macroscopic dynamics relative to the original source data.

#### Characterising the Emergent Dynamical Structure

To discover emergent macroscopic variables, or *n*-macros, we performed dimensionality reduction via dynamical independence (DI), using gradient descent to minimise dynamical dependence (DD) as defined in Eq. 3. For each subject and condition, 100 independent optimisations were conducted for each spatial scale 2≤*n*≤9, with each run starting from a random initialisation on the Stiefel manifold. Optimisation proceeded for up to 10, 000 iterations or until convergence was achieved to a tolerance below 10^−10^, using an adaptive step size. This process included a pre-optimisation phase based on the computationally-efficient proxy *F* ^*^(***X***→*M* ***X***), followed by refinement using the full spectral-domain DD functional *F* (***X***→*M* ***X***) [28]. Frequency integration was performed numerically from 0 Hz to the Nyquist frequency.

To characterise the *emergent dynamical structure* induced by the similarity of emergent *n*-macros, we computed pairwise *principal angles* between all optimised subspaces at each scale. This was implemented in MATLAB using the custom routine `subspacea.m`. The resulting 100×100 matrix of pairwise subspace distances represents the geometry of the solution space—how similar or distinct the *n*-macros are across independent optimisation runs.

Formally, for two *n*-dimensional subspaces of ℝ^*N*^, the *principal angles θ*_1_, …, *θ*_*n*_ ∈ [0, *π*/2] define an invariant metric on the Grassmannian manifold [119, 122]. The canonical metric between two subspaces is:

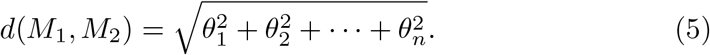

To compare subspaces across scales and subjects, we normalised this metric to lie in [0, 1], where 0 corresponds to perfect subspace alignment (identical *n*-macros), and 1 indicates orthogonality. For instance, when *n* = 2, this becomes:

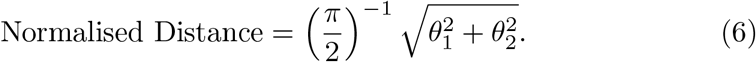

This *subspace geometry* captures the stability and consistency of the 100 DD-optimised *n*-macros. A tight clustering of angles around zero indicates the presence of a dominant macro-scale structure consistently identified across runs: i.e., a large basin of attraction around a local minimum in the optimisation landscape. In contrast, a broad distribution of angles suggests a more heterogeneous or fragmented dynamical organisation, potentially reflecting the absence of a unique emergent structure at that scale or condition.

#### Characterising Source-level Contributions to *n*-macros

To localise the contribution of individual sources to each emergent *n*-macro, we computed the angle between each source coordinate axis (in the original *N* = 46-dimensional source space) and the subspace defined by the *n*-macro. This projection-based approach yields a spatial interpretation of the macroscopic variable by quantifying how strongly each region contributes to the emergent dynamical structure—what we refer to as the *regional embedding*, or spatial localisation of the *n*-macro.

To assess whether a given regional contribution is statistically meaningful, we compared the observed projection against a null distribution derived from random subspaces. Under the null hypothesis, the *n*-macro is assumed to be sampled uniformly at random from the Grassmannian manifold 𝒢_*N*_ (*n*)—the space of all *n*-dimensional subspaces of ℝ^*N*^. This leads to a well-known result in algebraic geometry: the squared cosine of the angle between a fixed unit vector (representing a region) and a randomly chosen subspace follows a Beta distribution [123].

Specifically, if *θ* denotes the angle between a 1-dim subspace (i.e., a line, here the coordinate axis in ℝ^*n*^ corresponding to a source region) and a random *n*-dimensional subspace, then cos^2^(*θ*) follows a Beta distribution:

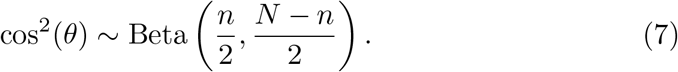

This distribution arises from a ratio of *χ*^2^ variables and captures the probability of a given region appearing to contribute to a subspace purely by chance. By comparing the observed squared cosine value for each source against this null Beta distribution, we compute a standardised effect size indicating whether the source contributes more strongly than expected under random projection. This provides a principled, probability-theoretic method to assess the *structural contribution* of each region to an emergent macroscopic process.

### 5.6 Statistical Comparison of Dynamical Dependence Differences: Degree of emergence

To statistically evaluate whether the degree of emergence—quantified by dynamical dependence—differed between wakefulness and anaesthetic conditions across spatial scales, we adopted a *participant-by-participant approach* using the Mann-Whitney U (MWU) test. This non-parametric two-tailed test (also known as the Wilcoxon rank-sum test) compares unpaired samples drawn from two populations—in our case DD values for the same subject under two conditions–against a null hypothesis that neither population stochastically dominates the other [62]. In contrast to an (unpaired) t-test, it avoids assumptions of normality.

For each participant *i*, and at each spatial scale *n*, the DD values from the wakefulness condition (sample size *n*_1_) were compared to those from the anaesthetic condition (sample size *n*_2_) using the MWU test, yielding a participant-specific test statistic *U*_*i*_.

The MWU statistic was then converted to a standardised Z-score using:

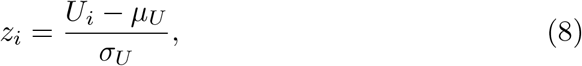

where the mean and standard deviation of the null distribution are given by:

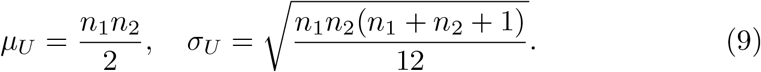

Here, *n*_1_ and *n*_2_ are the number of DD values (i.e., optimisation runs) in the wake and anaesthetic conditions respectively. The Z-score *z*_*i*_ captures the direction and magnitude of stochastic dominance for participant *i* at a given scale.

To derive a *group-level test statistic*, we aggregated these individual Z-scores across all *N* participants in a condition group (propofol, xenon, or ketamine) vs the wake condition as:

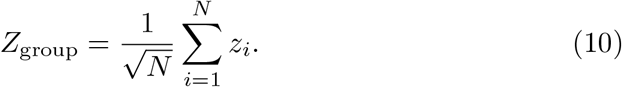

This yields a normally distributed variable under the null hypothesis that no consistent difference exists between conditions across participants. While this approach is not identical to Stouffer’s method, it is mathematically equivalent, as both rely on aggregating independent standard normal variables for group-level inference. The associated group-level p-value is then calculated as:

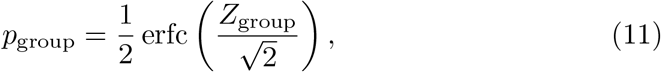

where erfc is the complementary error function. This provides a closed-form mapping from the group-level Z-score to a p-value under the standard normal distribution.

Finally, to correct for *multiple comparisons* across tested macroscales *n*, and conditions, we applied the Benjamini-Hochberg False Discovery Rate (FDR) procedure at a threshold of α = 0.05, ensuring statistical control of false positives across spatial scales.

This procedure enables robust, distribution-free testing of condition-related shifts in DD across participants and spatial scales, directly capturing stochastic dominance in the emergence measure.

### 5.7 Statistical Comparison of Similarity-Matrix Differences: Emergent dynamical structure

To assess how anaesthesia influences the emergence of structured dynamics across macroscopic levels of description, we analysed pairwise similarity matrices derived from optimised *n*-macros of high-density EEG time series. These similarity matrices were obtained for each subject, condition, and macroscopic spatial scale using the method outlined in preceding Section.

To summarise, we computed similarity matrices of shape 100×100 for each of eight macroscales (ranging from 2 to 9) and for two conditions: wakefulness and anaesthesia. For each matrix, we extracted the upper triangular values and computed a Structure-Fragmentation index of the emergent dynamics: the proportion of similarity values less than 0.0001. This measure reflects the organisation/fragmentation of the emergent landscape indicated by the resulting similarity matrix, with a higher proportion of near-zero values interpreted as reflecting greater structural similarity among the emergent macroscopic variables.

This SF index yields, for each subject and condition, a vector of 8 scalar values—one per macroscale level. We performed a repeated-measures analysis of variance (rmANOVA) separately for each anaesthetic agent (ketamine, propofol, xenon), with fixed effects of macro (8 levels) and condition (Wake, Anaesthesia), and subject included as a random effect. Partial eta-squared (*η*^2^) was reported as a measure of effect size.

To determine the directionality of any condition effects at specific macroscales, we conducted paired Wilcoxon signed-rank tests comparing wakefulness and anaesthesia for each macro level. The resulting *p*-values were corrected for multiple comparisons using the Benjamini-Hochberg false discovery rate (FDR) procedure. Significant macro levels were identified based on q-values (FDR-corrected *p*-values) below 0.05.

Visualisation of results included subject-level paired-line plots and overlaid group means across macro levels, with asterisks indicating macros where post hoc comparisons reached statistical significance. All statistical analyses were performed in Python using the `pingouin`, `scipy`, and `statsmodels` libraries.

### 5.8 Statistical Comparison of *β*-value Differences: Source-level contributions

To quantify the contribution of individual brain regions to emergent macroscale dynamics, we computed the hyperplane axis-angle *β*-statistic for each region, for each subject, condition, and macroscale dimensionality (*n*-macro size). To summarise, the *β*-statistic corresponds to the squared cosine of the angle between the axis of each region and the optimised hyperplane representing the macroscale variable. Intuitively, larger *β* values indicate stronger alignment (and thus stronger contribution) of that region to the *n*-macro.

Under the null hypothesis of random orientation of the hyperplane (no preferential contribution of any region), the *β*-statistic follows a known Beta(*m/*2, (*n* − *m*)*/*2) distribution, where *n* is the full system dimension (number of regions) and *m* is the dimension of the *n*-macro. Statistical significance was assessed by computing left-and right-tailed critical values from this null distribution. A region was considered to show significant participation if its *β* exceeded the right-tail threshold (close alignment with the macro), and significant non-participation if below the left-tail threshold.

To control for multiple comparisons across regions, a Bonferroni correction was applied to the significance level. This procedure was repeated for each subject, condition (wake, propofol, ketamine, xenon), and macroscale dimension (2 ≤ *m* ≤ 9). The resulting significant regions were visualised on the cortical surface (fsaverage, HCP-MMP1 parcellation, in the MNE-Python package) to generate the participant-level maps shown in Supplementary Sections 5–7.

On a group level, to analyse differences in the regional contribution *between* wake and anaesthetic conditions, we employed a two-level statistical approach. For each anaesthetic condition (propofol, xenon, and ketamine), we first performed paired Wilcoxon signed-rank tests to compare the *β*-statistics between wake and anaesthetic states. To assess the overall effect across all anaesthetic conditions, we utilised Stouffer’s Z-score method to aggregate the individual p-values^2^. In Stouffer’s method, individual p-values are transformed to Z-scores and combined as 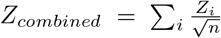, where *n* is the number of tests. The resulting Z-score follows a standard normal distribution, allowing for straightforward interpretation of the overall significance.

## Supplementary information

Supplementary Figures 1–3 and 5–7 provide additional analyses supporting the main findings. These include power spectral density (PSD) estimates across all conditions, grand-average pairwise-conditional Granger causality matrices, and statistical comparisons of *n*-macro similarity distributions using Kolmogorov–Smirnov (KS) tests (Supplementary Figures 1–3).

In addition, Supplementary Figures 5–7 present participant-level cortical maps of significant source contributions to the optimised *n*-macros for each anaesthetic condition (Propofol, Ketamine, Xenon), providing a spatially embedded perspective on how emergent macroscale structure might significantly reorganise under anaesthesia.

The full Supplementary Information is provided as a single PDF file.

## Acknowledgments

This work was supported in part by computational resources provided by the University of Melbourne’s SPARTAN HPC cluster. The authors thank members of the Human Experience Lab (HEx Lab) at the University of Melbourne, the DreamTeam at the Paris Brain Institute, and the Sussex Centre for Consciousness Science for their support and insights throughout the project.

## Declarations

### Funding

This work was supported by the European Research Council (ERC-StG SleepingAwake 101116748 to T.A.), the National Foundation for Medical Research and Innovation (APP1183280 to O.C. and B.M.), and internal research funding from the University of Melbourne (to O.C.). The work was also supported in part by the European Research Council (ERC) under the Horizon 2020 programme (Grant 101019254 to A.K.S. and L.B.).B.M. was further supported by the Australian Government Research Training Program Scholarship. The funders had no role in study design, data collection and analysis, decision to publish, or preparation of the manuscript.

### Conflict of interest/Competing interests

A.K.S. is an advisor to Conscium Ltd and to Alljoined Inc. All other authors declare no competing interests.

### Ethics approval

Ethical approval for the original study was obtained by the research team that collected the data [45], from the local ethics committee of the University of Liège, Belgium. The current analysis used only the publicly available, de-identified dataset and did not require additional ethical approval.

### Consent to participate

All participants provided written informed consent for the original study conducted by [45].

### Consent for publication

Not applicable. This study uses de-identified, previously published open-access data.

### Availability of data and materials

The EEG dataset analysed in this study is available in the open-access repository referenced in [45]. All relevant scripts, including those for preprocessing, source reconstruction, and dynamical independence analysis, are publicly available via the repositories listed below.

### Code availability

Custom methods and analysis used in this study are available in the GitHub repository [DI EEG] developed and maintained by B.M.: https://github.com/bmilinkovic/di_eeg.

Dynamical Independence (DI) analysis was performed using the SSDI toolbox developed by Lionel Barnett: https://github.com/lcbarnett/ssdi.

This toolbox depends on the MVGC2 toolbox for Granger causality analysis: https://github.com/lcbarnett/MVGC2.

### Authors’ contributions

B.M. conceived the study, performed the analysis, developed the software, and wrote the manuscript. T.A. and O.C. contributed to conceptual development, interpretation of results, and manuscript editing. L.B. developed the original theoretical framework and software for DI and assisted with methodological integration. A.K.S. provided guidance on theoretical framing and interpretation and contributed to manuscript development and editing. All authors reviewed and approved the final manuscript.

## Supplementary Information

### Supplementary Figures

#### Supplementary Section 1: Power Spectral Analysis

We examined the grand-average power spectral density (PSD) across conditions (Supplementary Fig. 1). The results replicate prior findings, including an increase in alpha power (8–13 Hz) under propofol compared to wakefulness [1, 2], a shift toward theta frequencies under ketamine [1], and a flattening of oscillatory peaks in xenon [3]. These spectral characteristics are consistent with previous analyses using the same dataset [4, 5].

**Supplementary Figure S1.**
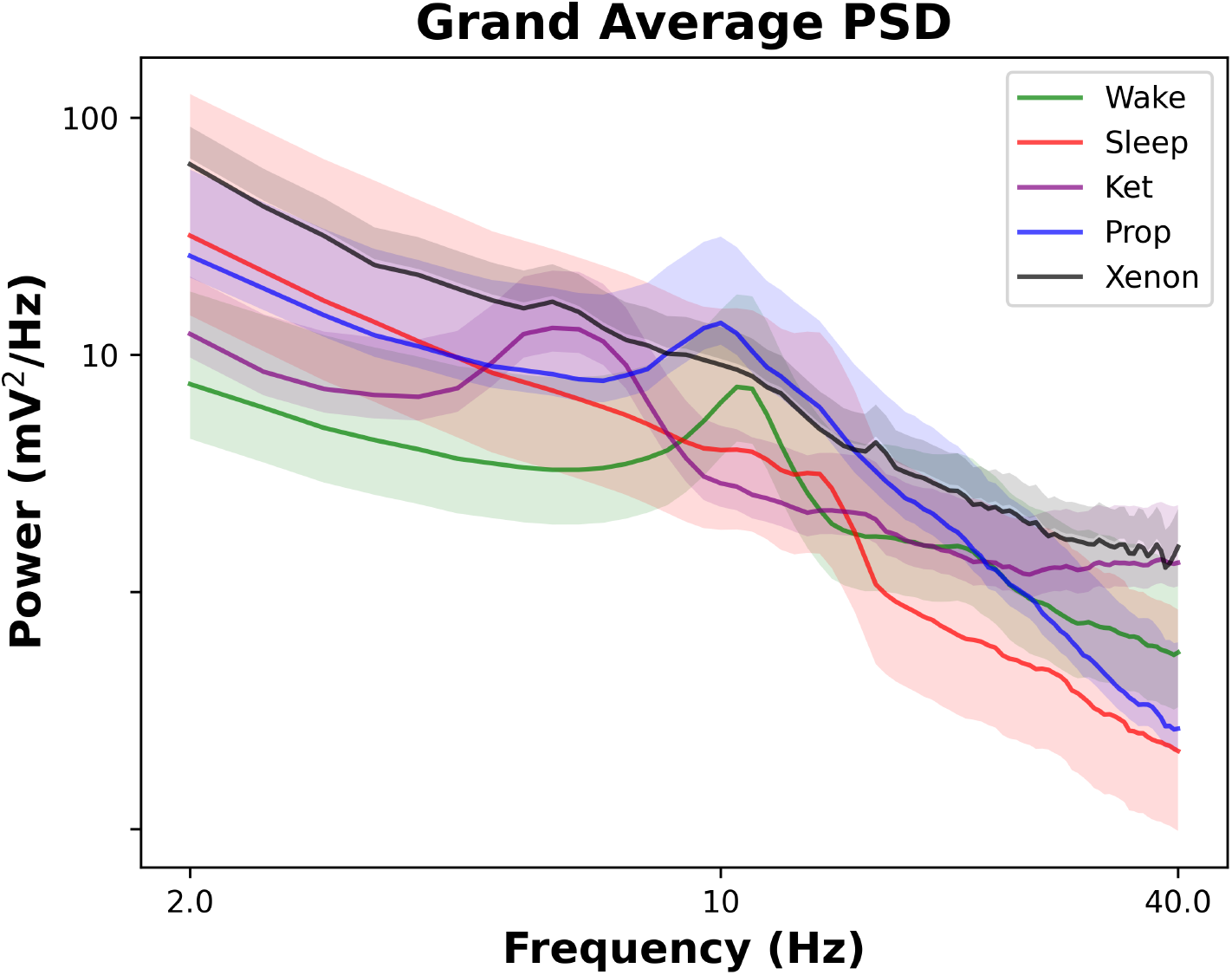
Grand-average Power Spectral Density across conditions. PSD estimates with 95% confidence intervals computed from 1,000 bootstrap samples. Ketamine shows a shift of the alpha peak toward lower frequencies, xenon flattens spectral peaks, and propofol increases alpha power.

#### Supplementary Section 2: Granger Causal Connectivity Matrices

We present grand-average pairwise conditional Granger causality (GC) matrices computed across all source-reconstructed regions for each condition (Supplementary Fig. 2). These matrices capture directed functional connectivity based on pairwise-conditional GC estimates.

**Supplementary Figure S2.**
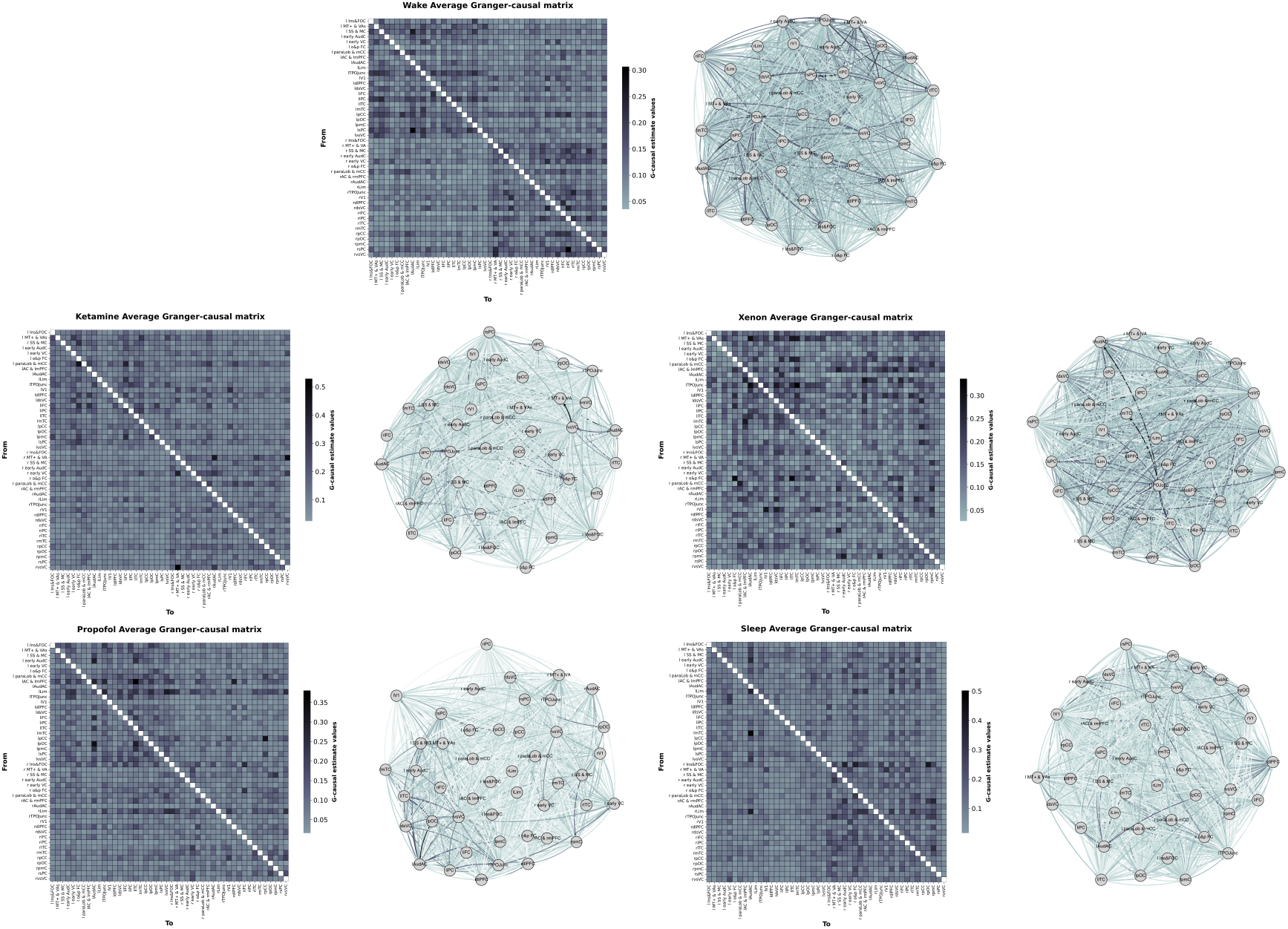
Granger Causal Connectivity Matrices. Grand-average pairwise conditional Granger-causality adjacency matrices for all conditions, computed from source-reconstructed EEG using the same regional atlas as shown in the main text.

#### Supplementary Section 3: Kolmogorov–Smirnov (KS) Tests on *n*-Macro Similarity Distributions

We assessed the distribution of similarity values between emergent *n*-macros across optimisation runs and conditions using Kolmogorov–Smirnov (KS) tests (Supplementary Fig. 3). KS statistics compared the empirical cumulative distribution functions (CDFs) of wakefulness against each anaesthetic condition. All conditions yielded significant differences after Bonferroni correction. propofol (*D* = 0.39, *p <* 0.0001), xenon (*D* = 0.28, *p <* 0.0001), and ketamine (*D* = 0.11, *p <* 0.001), indicating meaningful alterations in emergent dynamical structure across states.

**Supplementary Figure S3.**
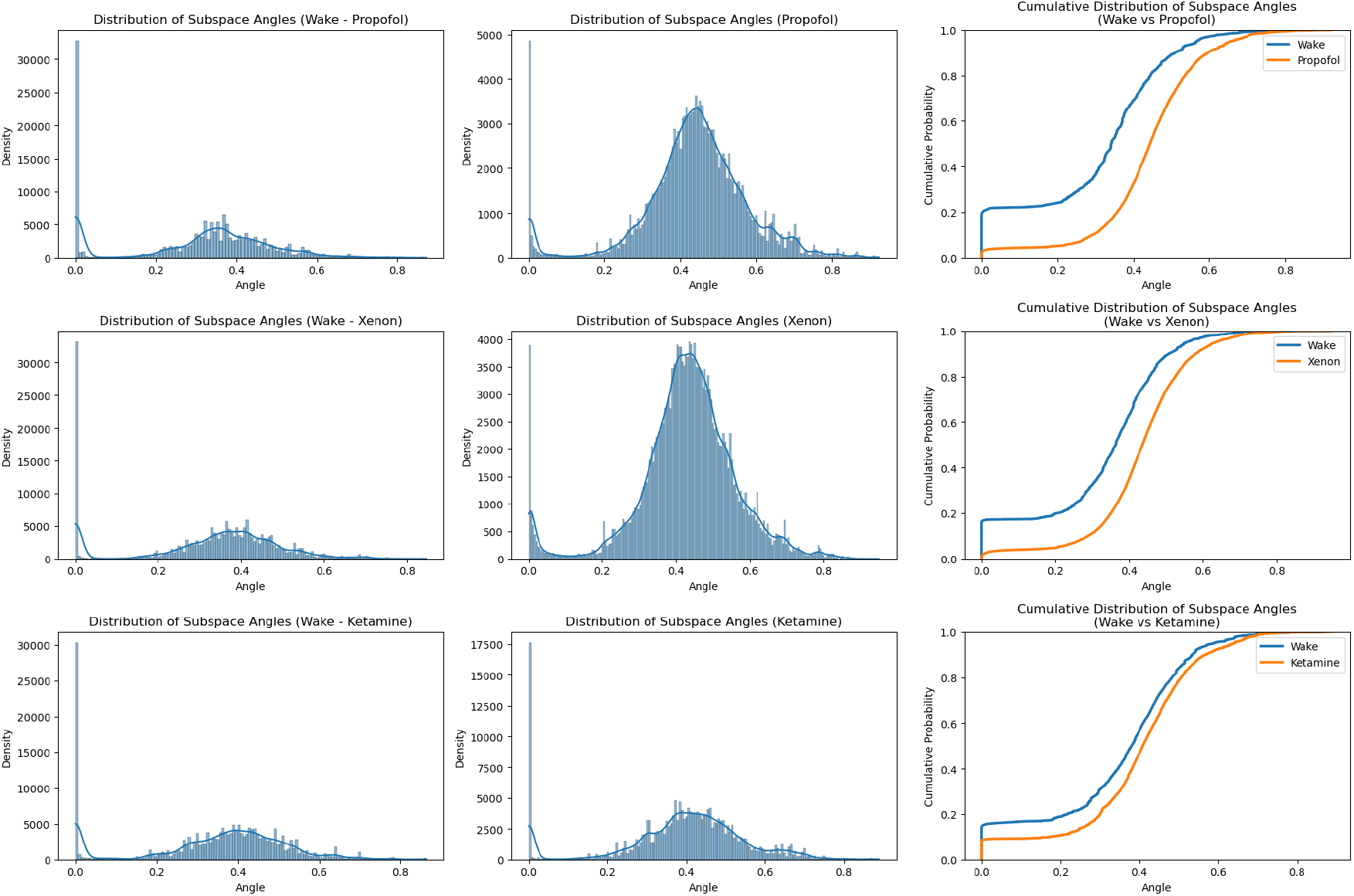
Distributions of n-Macro Similarity and KS Test Results. Empirical CDFs of n-macro similarity scores for wakefulness and each anaesthetic condition. KS tests reveal significantly different distributions between wakefulness and all anaesthetic conditions, indicating disrupted or fragmented macroscopic structure.

#### Supplementary Section 4: Explanation of Subject-Level Regional Contribution Maps of Significant Regional Contributions

To complement the group-level statistical analyses presented in the main text, we provide here individual participant-level maps of significant regional contributions to the optimised macroscale variables (*n*-macros) across Wake and each anaesthetic condition (Propofol, Ketamine, Xenon). These maps are intended to give a more detailed spatial perspective on how emergent macroscale structure reorganises under each anaesthetic, and to illustrate the degree of within-subject and between-subject variability across macroscales.

For each subject, we show results across macroscales 2–9, as well as an aggregate union map summarising regions that were significant at any scale. Regions are colour-coded to indicate whether they show significant participation in Wake, in the anaesthetic condition, or in both. These complementary visualisations provide an intuitive view of the spatial patterns underlying the quantitative measures reported in the main figures.

#### Supplementary Section 5: Wake vs Propofol — Significant Regional Contributions

In this section we provide individual participant-level maps showing the significant cortical regions contributing to the optimised macroscale variables (*n*-macros) for Wake vs Propofol. For each participant and each macroscale (2–9), regions are colour-coded according to whether they show significant participation in Wake only, Propofol only, or both. An aggregate union map (“Any Macroscale”) shows regions that were significant at any tested scale. These maps provide a spatial perspective on how macroscale participation patterns reorganise under propofol anaesthesia.

#### Supplementary Section 6: Wake vs Xenon — Significant Regional Contributions

Here we display participant-level maps for Wake vs Xenon, again showing significant regional participation in the optimised macroscales across individual subjects. The union map highlights regions consistently participating at any scale. These results reveal widespread and often bilateral changes in macroscale structure under xenon anaesthesia.

#### Supplementary Section 7: Wake vs Ketamine — Significant Regional Contributions

This section presents individual participant-level maps for Wake vs Ketamine. The same conventions are used as for the previous figure: regions significantly contributing to the optimised macroscales are displayed for each participant and each scale, with a union map summarising robust contributions across scales. These maps illustrate the heterogeneous and spatially distributed changes in macroscale participation patterns under ketamine.

**Supplementary Figure S4.**
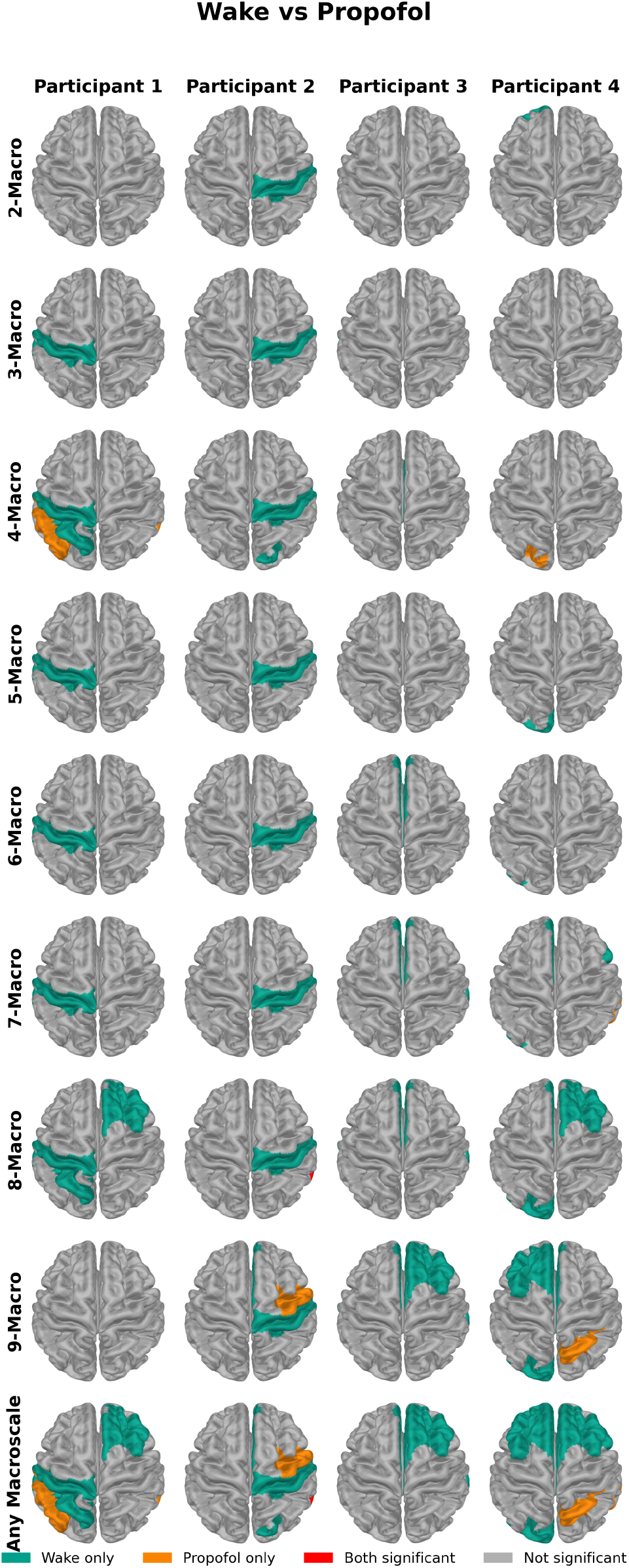
Wake vs Propofol: significant regional contributions to optimised macroscales (2–9) and union map (“Any Macroscale”) for each participant (N=4). Green = Wake only; Orange = Propofol only; Red = both; Grey = non-significant.

**Supplementary Figure S6.**
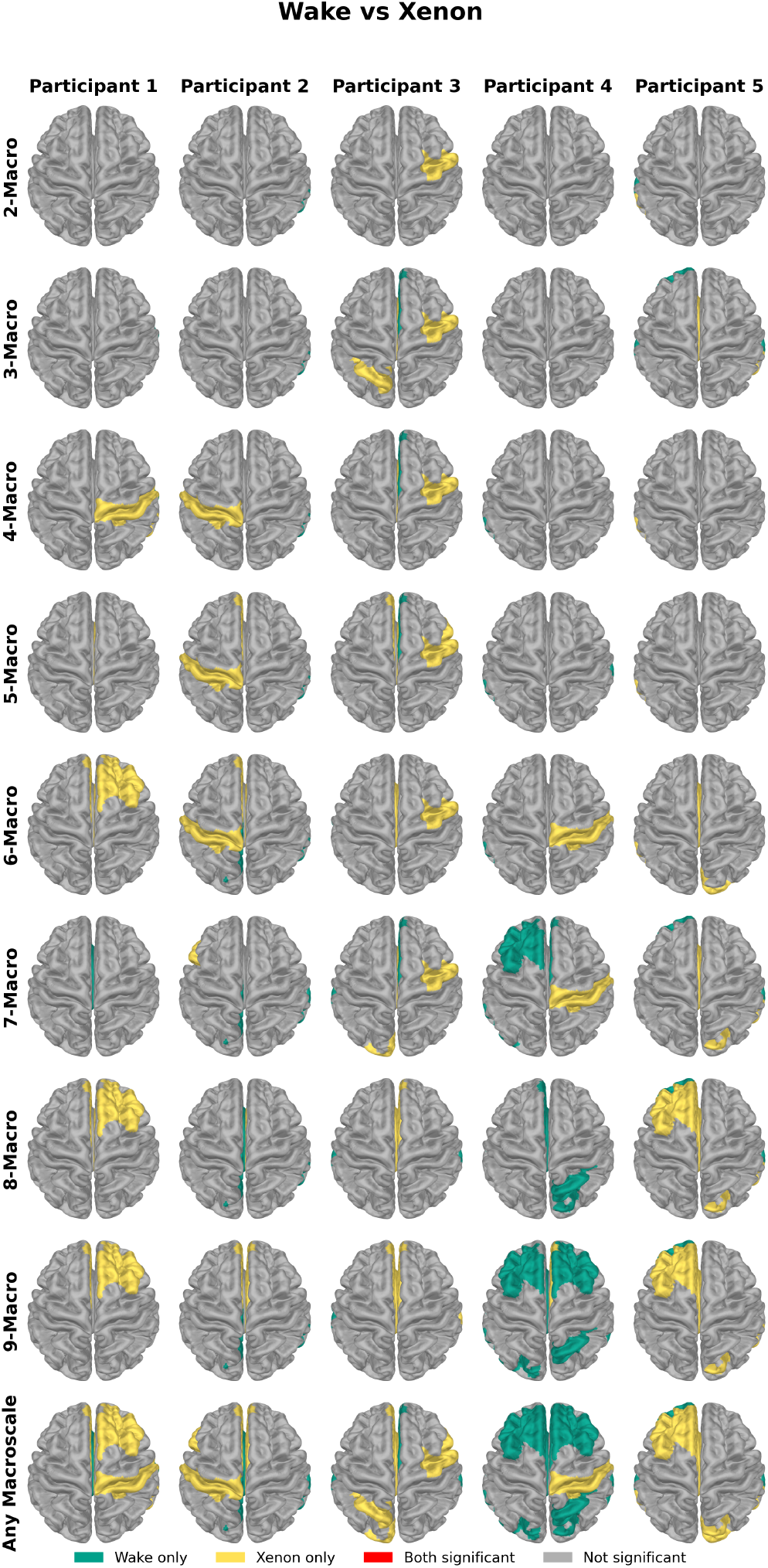
Wake vs Xenon: significant regional contributions to optimised macroscales (2–9) and union map (“Any Macroscale”) for each participant (N=5). Green = Wake only; Yellow = Xenon only; Red = both; Grey = non-significant.

**Supplementary Figure S7.**
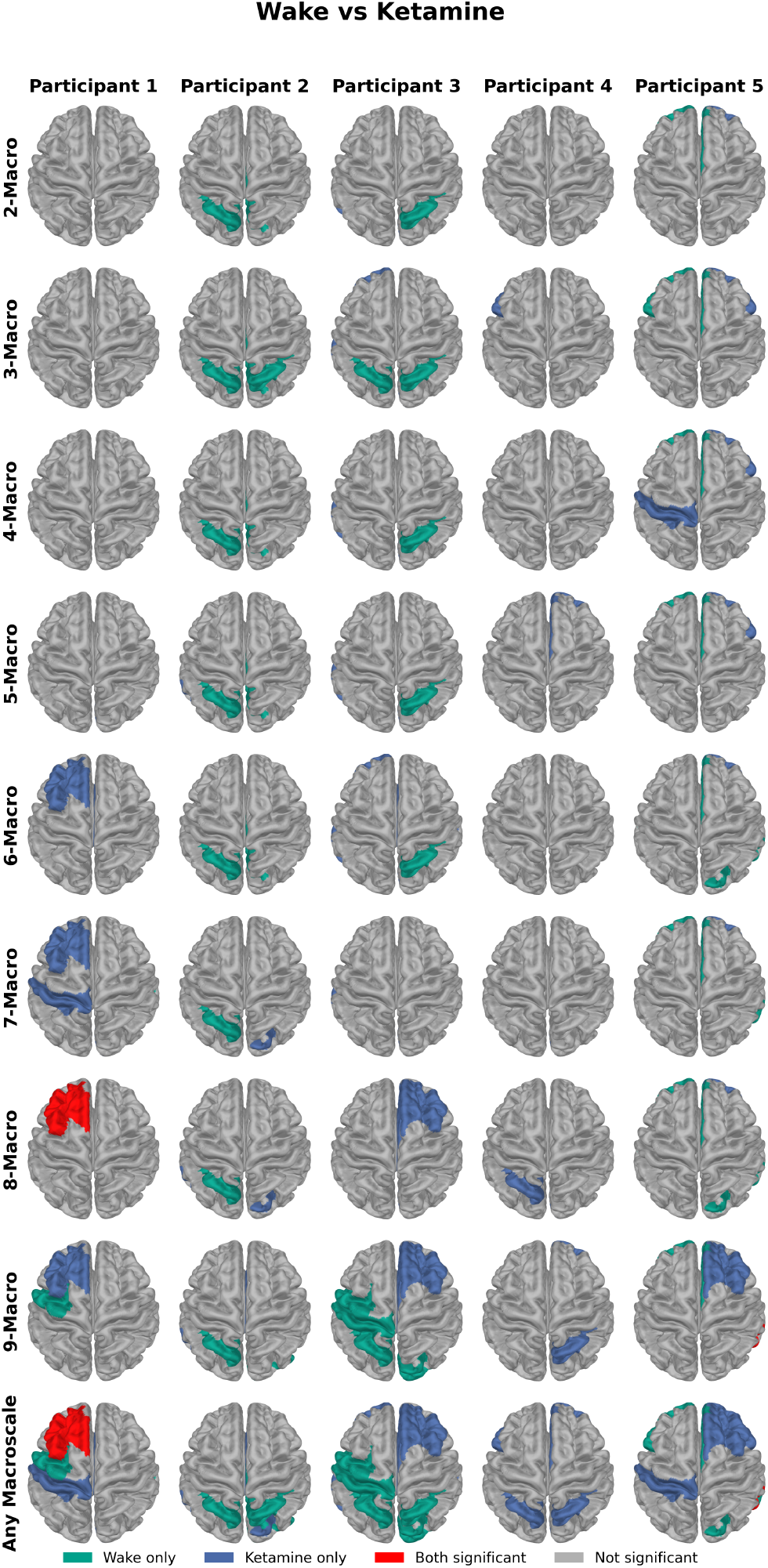
Wake vs Ketamine: significant regional contributions to optimised macroscales (2–9) and union map (“Any Macroscale”) for each participant (N=5). Green = Wake only; Blue = Ketamine only; Red = both; Grey = non-significant.

## Footnotes

1 As mentioned, DD is well-defined as a real-valued function on the associated Grassmannian manifold 𝒢_*N*_ (*n*). As a ‘cost function’ for optimisation, DD is in general highly multi-modal; as a consequence, as we shall see below, the optimisation process deployed in this paper will frequently find *n*-macros with DD a local, rather than global minimum on 𝒢_*N*_ (*n*).

2 Stouffer’s method was chosen over alternatives such as Fisher’s method because it is optimally suited for detecting consistent effects across multiple conditions, rather than being overly sensitive to strong effects in a single condition.

## Notes

https://github.com/bmilinkovic/di_eeg

## References

[1] Friston, K. Hierarchical models in the brain. PLoS computational biology 4 (11), e1000211 (2008).

[2] Raut, R. V., Snyder, A. Z. & Raichle, M. E. Hierarchical dynamics as a macroscopic organizing principle of the human brain. Proceedings of the National Academy of Sciences 117 (34), 20890–20897 (2020).

[3] Deco, G., Vidaurre, D. & Kringelbach, M. L. Revisiting the global workspace orchestrating the hierarchical organization of the human brain. Nature human behaviour 5 (4), 497–511 (2021).

[4] Kringelbach, M. L., Perl, Y. S., Tagliazucchi, E. & Deco, G. Toward naturalistic neuroscience: Mechanisms underlying the flattening of brain hierarchy in movie-watching compared to rest and task. Science Advances 9 (2), eade6049 (2023).

[5] Li, A. et al. Hierarchical fluctuation shapes a dynamic flow linked to states of consciousness. Nature communications 14 (1), 3238 (2023).

[6] Luppi, A. I. et al. Reduced emergent character of neural dynamics in patients with a disrupted connectome. Neuroimage 269, 119926 (2023).

[7] Kelso, J. S. The dynamic brain in action: Coordinative structures, criticality, and coordination dynamics. Criticality in neural systems 67–104 (2014).

[8] Breakspear, M. Dynamic models of large-scale brain activity. Nature neuroscience 20 (3), 340–352 (2017).

[9] Rubinov, M. & Sporns, O. Complex network measures of brain connectivity: uses and interpretations. Neuroimage 52 (3), 1059–1069 (2010).

[10] Demertzi, A., Soddu, A. & Laureys, S. Consciousness supporting networks. Current opinion in neurobiology 23 (2), 239–244 (2013).

[11] Uehara, T. et al. Efficiency of a “small-world” brain network depends on consciousness level: a resting-state fmri study. Cerebral cortex 24 (6), 1529–1539 (2014).

[12] Lee, H. et al. Relationship of critical dynamics, functional connectivity, and states of consciousness in large-scale human brain networks. Neuroimage 188, 228–238 (2019).

[13] Fornito, A., Zalesky, A. & Breakspear, M. The connectomics of brain disorders. Nature Reviews Neuroscience 16 (3), 159–172 (2015).

[14] Fornito, A., Zalesky, A. & Bullmore, E. Fundamentals of brain network analysis (Academic press, 2016).

[15] Bassett, D. S. & Sporns, O. Network neuroscience. Nature neuroscience 20 (3), 353–364 (2017).

[16] Avena-Koenigsberger, A., Misic, B. & Sporns, O. Communication dynamics in complex brain networks. Nature reviews neuroscience 19 (1), 17–33 (2018).

[17] Battiston, F. et al. Networks beyond pairwise interactions: Structure and dynamics. Physics Reports 874, 1–92 (2020).

[18] Battiston, F. & Petri, G. Higher-Order Systems (Springer, 2022).

[19] Cabral, J. et al. Metastable oscillatory modes emerge from synchronization in the brain spacetime connectome. Communications Physics 5 (1), 184 (2022).

[20] Cabral, J., Fernandes, F. F. & Shemesh, N. Intrinsic macroscale oscillatory modes driving long range functional connectivity in female rat brains detected by ultrafast fmri. Nature Communications 14 (1), 375 (2023).

[21] Atasoy, S., Deco, G., Kringelbach, M. L. & Pearson, J. Harmonic brain modes: a unifying framework for linking space and time in brain dynamics. The Neuroscientist 24 (3), 277–293 (2018).

[22] Luppi, A. I. et al. Connectome harmonic decomposition of human brain dynamics reveals a landscape of consciousness. BioRxiv (2020).

[23] Tononi, G., Edelman, G. M. & Sporns, O. Complexity and coherency: integrating information in the brain. Trends in cognitive sciences 2 (12), 474–484 (1998).

[24] Seth, A. K. & Bayne, T. Theories of consciousness. Nature Reviews Neuroscience 23 (7), 439–452 (2022).

[25] Atmanspacher, H. & Bishop, R. C. Stability conditions in contextual emergence. Chaos and Complexity Letters 2 (2/3), 139–150 (2007).

[26] Hoel, E. P., Albantakis, L. & Tononi, G. Quantifying causal emergence shows that macro can beat micro. Proceedings of the National Academy of Sciences 110 (49), 19790–19795 (2013).

[27] Rosas, F. E. et al. Reconciling emergences: An information-theoretic approach to identify causal emergence in multivariate data. PLoS computational biology 16 (12), e1008289 (2020).

[28] Barnett, L. & Seth, A. K. Dynamical independence: discovering emergent macroscopic processes in complex dynamical systems. Physical Review E 108 (1), 014304 (2023).

[29] Allefeld, C., Atmanspacher, H. & Wackermann, J. Mental states as macrostates emerging from brain electrical dynamics. Chaos: An Interdisciplinary Journal of Nonlinear Science 19 (1), 015102 (2009).

[30] Shah, P. et al. Characterizing the role of the structural connectome in seizure dynamics. Brain 142 (7), 1955–1972 (2019).

[31] Luppi, A. I. et al. A synergistic workspace for human consciousness revealed by integrated information decomposition. BioRxiv 2020–11 (2020).

[32] Luppi, A. I. et al. Reduced emergent character of neural dynamics in patients with a disrupted connectome. bioRxiv (2022).

[33] Herzog, R. et al. Genuine high-order interactions in brain networks and neurodegeneration. Neurobiology of Disease 175, 105918 (2022).

[34] Varley, T. F., Pope, M., Faskowitz, J. & Sporns, O. Multivariate information theory uncovers synergistic subsystems of the human cerebral cortex. Communications biology 6 (1), 451 (2023).

[35] Herzog, R. et al. High-order brain interactions in ketamine during rest and task: a double-blinded cross-over design using portable eeg on male participants. Translational Psychiatry 14 (1), 310 (2024).

[36] Luppi, A. I. et al. A synergistic core for human brain evolution and cognition. Nature Neuroscience 25 (6), 771–782 (2022).

[37] Luppi, A. I., Rosas, F. E., Mediano, P. A., Menon, D. K. & Stamatakis, E. A. Information decomposition and the informational architecture of the brain. Trends in Cognitive Sciences (2024).

[38] Milinkovic, B., Barnett, L., Carter, O., Seth, A. K. & Andrillon, T. Capturing the emergent dynamical structure in biophysical neural models. PLOS Computational Biology 21 (5), e1012572 (2025).

[39] Valli, K. et al. Subjective experiences during dexmedetomidine-or propofol-induced unresponsiveness and non-rapid eye movement sleep in healthy male subjects. British Journal of Anaesthesia 131 (2), 348–359 (2023).

[40] Krestow, M. The effect of post-anaesthetic dreaming on patient acceptance of ketamine anaesthesia: a comparison with thiopentone-nitrous oxide anaesthesia. Canadian Anaesthetists’ Society Journal 21, 385–389 (1974).

[41] Radek, L. et al. Subjective experiences are similar during anaesthetic-induced unresponsiveness and non-rapid eye movement sleep. British Journal of Anaesthesia 130 (2), e397–e398 (2023).

[42] Kantonen, O. et al. Decreased thalamic activity is a correlate for disconnectedness during anesthesia with propofol, dexmedetomidine and sevoflurane but not s-ketamine. Journal of Neuroscience 43 (26), 4884–4895 (2023).

[43] Eisen, A. J. et al. Propofol anesthesia destabilizes neural dynamics across cortex. Neuron (2024).

[44] Purdon, P. L., Sampson, A., Pavone, K. J. & Brown, E. N. Clinical electroencephalography for anesthesiologists: part i: background and basic signatures. Anesthesiology 123 (4), 937–960 (2015).

[45] Sarasso, S. et al. Consciousness and complexity during unresponsiveness induced by propofol, xenon, and ketamine. Current Biology 25 (23), 3099–3105 (2015).

[46] McGuigan, S. et al. Comparison of the spectral features of the frontal electroencephalogram in patients receiving xenon and sevoflurane general anesthesia. Anesthesia & Analgesia 133 (5), 1269–1279 (2021).

[47] Murphy, M. et al. Propofol anesthesia and sleep: a high-density eeg study. Sleep 34 (3), 283–291 (2011).

[48] Purdon, P. L. et al. Electroencephalogram signatures of loss and recovery of consciousness from propofol. Proceedings of the National Academy of Sciences 110 (12), E1142–E1151 (2013).

[49] Koskinen, M., Seppänen, T., Tuukkanen, J., Yli-Hankala, A. & Jäntti, V. Propofol anesthesia induces phase synchronization changes in eeg. Clinical neurophysiology 112 (2), 386–392 (2001).

[50] Johnson, B., Sleigh, J., Kirk, I. & Williams, M. High-density eeg mapping during general anaesthesia with xenon and propofol: a pilot study. Anaesthesia and intensive care 31 (2), 155–163 (2003).

[51] Franks, N. P. General anaesthesia: from molecular targets to neuronal pathways of sleep and arousal. Nature Reviews Neuroscience 9 (5), 370– 386 (2008).

[52] Isomura, Y. et al. Integration and segregation of activity in entorhinal-hippocampal subregions by neocortical slow oscillations. Neuron 52 (5), 871–882 (2006).

[53] Breshears, J. D. et al. Stable and dynamic cortical electrophysiology of induction and emergence with propofol anesthesia. Proceedings of the National Academy of Sciences 107 (49), 21170–21175 (2010).

[54] Massimini, M. et al. Sleep-like cortical dynamics during wakefulness and their network effects following brain injury. Nature Communications 15 (1), 7207 (2024).

[55] Hudetz, A. G. General anesthesia and human brain connectivity. Brain connectivity 2 (6), 291–302 (2012).

[56] Juel, B. E., Romundstad, L., Kolstad, F., Storm, J. F. & Larsson, P. G. Distinguishing anesthetized from awake state in patients: a new approach using one second segments of raw eeg. Frontiers in Human Neuroscience 12, 40 (2018).

[57] Casali, A. G. et al. A theoretically based index of consciousness independent of sensory processing and behavior. Science translational medicine 5 (198), 198ra105–198ra105 (2013).

[58] Schartner, M. et al. Complexity of multi-dimensional spontaneous eeg decreases during propofol induced general anaesthesia. PloS one 10 (8), e0133532 (2015).

[59] Colombo, M. A. et al. The spectral exponent of the resting eeg indexes the presence of consciousness during unresponsiveness induced by propofol, xenon, and ketamine. NeuroImage 189, 631–644 (2019).

[60] Maschke, C., Duclos, C., Owen, A. M., Jerbi, K. & Blain-Moraes, S. Aperiodic brain activity and response to anesthesia vary in disorders of consciousness. NeuroImage 275, 120154 (2023).

[61] Varley, T. F., Sporns, O., Puce, A. & Beggs, J. Differential effects of propofol and ketamine on critical brain dynamics. PLoS computational biology 16 (12), e1008418 (2020).

[62] Barrett, G. F. & Donald, S. G. Consistent tests for stochastic dominance. Econometrica 71 (1), 71–104 (2003).

[63] Whitlock, M. C. Combining probability from independent tests: the weighted z-method is superior to fisher’s approach. Journal of evolutionary biology 18 (5), 1368–1373 (2005).

[64] Cofré, R. & Destexhe, A. Entropy and complexity tools across scales in neuroscience: A review. Entropy 27 (2), 115 (2025).

[65] Schartner, M. M., Carhart-Harris, R. L., Barrett, A. B., Seth, A. K. & Muthukumaraswamy, S. D. Increased spontaneous meg signal diversity for psychoactive doses of ketamine, lsd and psilocybin. Scientific reports 7 (1), 46421 (2017).

[66] Kleiner, J. & Ludwig, T. If consciousness is dynamically relevant, artificial intelligence isn’t conscious. arXiv preprint 2304.05077 (2023).

[67] Aru, J., Suzuki, M. & Larkum, M. E. Cellular mechanisms of conscious processing. Trends in cognitive sciences 24 (10), 814–825 (2020).

[68] Aru, J., Larkum, M. E. & Shine, J. M. The feasibility of artificial consciousness through the lens of neuroscience. Trends in neurosciences 46 (12), 1008–1017 (2023).

[69] Seth, A. K. Conscious artificial intelligence and biological naturalism. Behavioral and Brain Sciences 1–42 (2024).

[70] Pessoa, L. The entangled brain. Journal of cognitive neuroscience 35 (3), 349–360 (2023).

[71] Goldman, J. S. et al. Bridging Single Neuron Dynamics to Global Brain States. Frontiers in Systems Neuroscience 0, 75 (2019).

[72] Di Volo, M. & Destexhe, A. Optimal responsiveness and information flow in networks of heterogeneous neurons. Scientific reports 11 (1), 17611 (2021).

[73] Barnett, L., Lizier, J. T., Harré, M., Seth, A. K. & Bossomaier, T. Information flow in a kinetic ising model peaks in the disordered phase. Physical review letters 111 (17), 177203 (2013).

[74] Pigorini, A. et al. Bistability breaks-off deterministic responses to intracortical stimulation during non-rem sleep. Neuroimage 112, 105–113 (2015).

[75] Achermann, P. & BorbÉly, A. Temporal evolution of coherence and power in the human sleep electroencephalogram. Journal of sleep research 7 (S1), 36–41 (1998).

[76] Massimini, M. et al. Breakdown of cortical effective connectivity during sleep. Science 309 (5744), 2228–2232 (2005).

[77] Siclari, F. et al. The neural correlates of dreaming. Nature neuroscience 20 (6), 872–878 (2017).

[78] Koch, C., Massimini, M., Boly, M. & Tononi, G. Neural correlates of consciousness: progress and problems. Nature reviews neuroscience 17 (5), 307–321 (2016).

[79] McCulloch, W. S. A heterarchy of values determined by the topology of nervous nets. The bulletin of mathematical biophysics 7, 89–93 (1945).

[80] Semedo, J. D., Zandvakili, A., Machens, C. K., Byron, M. Y. & Kohn, A. Cortical areas interact through a communication subspace. Neuron 102 (1), 249–259 (2019).

[81] McMillen, P. & Levin, M. Collective intelligence: A unifying concept for integrating biology across scales and substrates. Communications Biology 7 (1), 378 (2024).

[82] Ebitz, R. B. & Hayden, B. Y. The population doctrine in cognitive neuroscience. Neuron 109 (19), 3055–3068 (2021).

[83] Thibeault, V., Allard, A. & Desrosiers, P. The low-rank hypothesis of complex systems. Nature Physics 20 (2), 294–302 (2024).

[84] Gao, J. Intrinsic simplicity of complex systems. Nature Physics 20 (2), 184–185 (2024).

[85] Mediano, P. A. M. et al. Towards an extended taxonomy of information dynamics via integrated information decomposition. arXiv preprint 2109.13186 (2021).

[86] Williams, P. L. & Beer, R. D. Nonnegative decomposition of multivariate information. arXiv preprint 1004.2515 (2010).

[87] Timme, N., Alford, W., Flecker, B. & Beggs, J. M. Synergy, redundancy, and multivariate information measures: an experimentalist’s perspective. Journal of computational neuroscience 36, 119–140 (2014).

[88] Timme, N. M. & Lapish, C. A tutorial for information theory in neuroscience. eneuro 5 (3) (2018).

[89] Gatica, M. et al. High-order interdependencies in the aging brain. Brain connectivity 11 (9), 734–744 (2021).

[90] Varley, T. F., Pope, M., Grazia, M., Joshua & Sporns, O. Partial entropy decomposition reveals higher-order information structures in human brain activity. Proceedings of the National Academy of Sciences 120 (30), e2300888120 (2023).

[91] Stramaglia, S., Scagliarini, T., Daniels, B. C. & Marinazzo, D. Quantifying dynamical high-order interdependencies from the o-information: an application to neural spiking dynamics. Frontiers in Physiology 11, 595736 (2021).

[92] Sherrill, S. P., Timme, N. M., Beggs, J. M. & Newman, E. L. Correlated activity favors synergistic processing in local cortical networks in vitro at synaptically relevant timescales. Network Neuroscience 4 (3), 678–697 (2020).

[93] Jansma, A., Mediano, P. A. & Rosas, F. E. The fast m “ obius transform: An algebraic approach to information decomposition. arXiv preprint 2410.06224 (2024).

[94] Mediano, P. A. et al. Greater than the parts: a review of the information decomposition approach to causal emergence. Philosophical Transactions of the Royal Society A 380 (2227), 20210246 (2022).

[95] Luppi, A. I. et al. What it is like to be a bit: an integrated information decomposition account of emergent mental phenomena. Neuroscience of consciousness 2021 (2), niab027 (2021).

[96] Humphries, M. D. Strong and weak principles of neural dimension reduction. arXiv preprint 2011.08088 (2020).

[97] Griebenow, R., Klein, B. & Hoel, E. Finding the right scale of a network: efficient identification of causal emergence through spectral clustering. arXiv preprint 1908.07565 (2019).

[98] Klein, B. & Hoel, E. The emergence of informative higher scales in complex networks. Complexity 2020, 1–12 (2020).

[99] Saggar, M. et al. Towards a new approach to reveal dynamical organization of the brain using topological data analysis. Nature communications 9 (1), 1399 (2018).

[100] Rosas, F. E., Mediano, P. A., Gastpar, M. & Jensen, H. J. Quantifying high-order interdependencies via multivariate extensions of the mutual information. Physical Review E 100 (3), 032305 (2019).

[101] Albantakis, L. et al. Integrated information theory (iit) 4.0: formulating the properties of phenomenal existence in physical terms. PLoS computational biology 19 (10), e1011465 (2023).

[102] Boly, M. et al. Are the neural correlates of consciousness in the front or in the back of the cerebral cortex? clinical and neuroimaging evidence. Journal of Neuroscience 37 (40), 9603–9613 (2017).

[103] Dehaene, S., Kerszberg, M. & Changeux, J.-P. A neuronal model of a global workspace in effortful cognitive tasks. Proceedings of the national Academy of Sciences 95 (24), 14529–14534 (1998).

[104] Dehaene, S. & Changeux, J.-P. Experimental and theoretical approaches to conscious processing. Neuron 70 (2), 200–227 (2011).

[105] Melloni, L., Mudrik, L., Pitts, M. & Koch, C. Making the hard problem of consciousness easier. Science 372 (6545), 911–912 (2021).

[106] Consortium, C. et al. An adversarial collaboration to critically evaluate theories of consciousness. BioRxiv 2023–06 (2023).

[107] Barnett, L., Barrett, A. B. & Seth, A. K. Granger causality and transfer entropy are equivalent for gaussian variables. Physical review letters 103 (23), 238701 (2009).

[108] Barnett, L. & Seth, A. K. The mvgc multivariate granger causality toolbox: a new approach to granger-causal inference. Journal of neuroscience methods 223, 50–68 (2014).

[109] Barnett, L. & Seth, A. K. Granger causality for state-space models. Physical Review E 91 (4), 040101 (2015).

[110] Hannan, E. J. & Deistler, M. The statistical theory of linear systems (SIAM, 2012).

[111] Cunningham, J. P. & Yu, B. M. Dimensionality reduction for large-scale neural recordings. Nature neuroscience 17 (11), 1500–1509 (2014).

[112] Burges, C. J. Dimension reduction: A guided tour. Foundations and Trends® in Machine Learning 2 (4), 275–365 (2010).

[113] Pang, R., Lansdell, B. J. & Fairhall, A. L. Dimensionality reduction in neuroscience. Current Biology 26 (14), R656–R660 (2016).

[114] Pearson, K. Liii. on lines and planes of closest fit to systems of points in space. The London, Edinburgh, and Dublin philosophical magazine and journal of science 2 (11), 559–572 (1901).

[115] Bishop, C. M. & Nasrabadi, N. M. Pattern recognition and machine learning Vol. 4 (Springer, 2006).

[116] Switzer, P. Min/max autocor-relation factors for multivariate spatial imagery. Technical report, Dept. of Statistics, Stanford University (1984)

[117] Churchland, M. M. et al. Neural population dynamics during reaching. Nature 487 (7405), 51–56 (2012).

[118] Bendokat, T., Zimmermann, R. & Absil, P.-A. A grassmann manifold handbook: Basic geometry and computational aspects. arXiv preprint 2011.13699 (2020).

[119] Absil, P.-A., Mahony, R. & Sepulchre, R. Optimization algorithms on matrix manifolds (Princeton University Press, 2008).

[120] Barnett, L., Barrett, A. B. & Seth, A. K. Granger causality and transfer entropy Are equivalent for gaussian variables. Physical Review Letters 103 (23) (2009).

[121] Barnett, L. & Bossomaier, T. Transfer Entropy as a Log-likelihood Ratio (2012).

[122] Wong, Y.-C. Differential geometry of grassmann manifolds. Proceedings of the National Academy of Sciences 57 (3), 589–594 (1967).

[123] Frankl, P. & Maehara, H. Some geometric applications of the beta distribution. Annals of the Institute of Statistical Mathematics 42, 463–474 (1990).

## References

[1] Patrick L Purdon, Eric T Pierce, Eran A Mukamel, Michael J Prerau, John L Walsh, Kin Foon K Wong, Andres F Salazar-Gomez, Priscilla G Harrell, Aaron L Sampson, Aylin Cimenser, et al. Electroencephalogram signatures of loss and recovery of consciousness from propofol. Proceedings of the National Academy of Sciences, 110(12):E1142–E1151, 2013.

[2] Miika Koskinen, Tapio Seppänen, Johanna Tuukkanen, Arvi Yli-Hankala, and Ville Jäntti. Propofol anesthesia induces phase synchronization changes in eeg. Clinical neurophysiology, 112(2):386–392, 2001.

[3] Steven McGuigan, Lisbeth Evered, Brendan Silbert, David A Scott, John R Cormack, Abarna Devapalasundaram, and David TJ Liley. Comparison of the spectral features of the frontal electroencephalogram in patients receiving xenon and sevoflurane general anesthesia. Anesthesia & Analgesia, 133(5):1269–1279, 2021.

[4] Simone Sarasso, Melanie Boly, Martino Napolitani, Olivia Gosseries, Vanessa Charland-Verville, Silvia Casarotto, Mario Rosanova, Adenauer Girardi Casali, Jean-Francois Brichant, and Pierre Boveroux. Consciousness and complexity during unresponsiveness induced by propofol, xenon, and ketamine. Current Biology, 25(23):3099–3105, 2015.

[5] Michele Angelo Colombo, Martino Napolitani, Melanie Boly, Olivia Gosseries, Silvia Casarotto, Mario Rosanova, Jean-Francois Brichant, Pierre Boveroux, Steffen Rex, and Steven Laureys. The spectral exponent of the resting eeg indexes the presence of consciousness during unresponsiveness induced by propofol, xenon, and ketamine. NeuroImage, 189:631–644, 2019.

